# Joint effects of social interactions and environmental challenges on physiology, internal microbiome, and reproductive performance in tree swallows (*Tachycineta bicolor*)

**DOI:** 10.1101/2023.01.05.522952

**Authors:** Conor C. Taff, Sabrina M. McNew, Cedric Zimmer, Jennifer J. Uehling, Jennifer L. Houtz, Thomas A. Ryan, David Chang van Oordt, Allison S. Injaian, Maren N. Vitousek

## Abstract

The social environment that individuals experience appears to be a particularly salient mediator of stress resilience, as the nature and valence of social interactions are often related to subsequent health, physiology, microbiota, and overall stress resilience. Relatively few studies have simultaneously manipulated the social environment and ecological challenges under natural conditions. Here, we report the results of experiments in wild tree swallows (*Tachycineta bicolor*) in which we manipulated both ecological challenges (predator encounters and flight efficiency reduction) and social interactions (by experimental dulling of a social signal). In two experiments conducted in separate years, we reversed the order of these treatments so that females experienced either an altered social environment followed by a challenge or vice-versa. Before, during, and after treatments were applied, we tracked breeding success, morphology and physiology (mass, corticosterone, and glucose), social interactions via an RFID sensor network, cloacal microbiome diversity, and fledging success. Overall, we found that predator exposure during the nestling period reduced the likelihood of fledging and that signal manipulation sometimes altered social interactions, but little evidence that the two categories of treatment interacted with each other. We discuss the implications of our results for understanding what types of challenges and what conditions are most likely to result in interactions between the social environment and ecological challenges.

## INTRODUCTION

The social environment can profoundly influence how animals cope with future challenges. Social position and social connectedness have emerged as key predictors of the psychophysiological impact of stress (Charuvastra & Cloitre, 2008; Holt-Lunstad et al., 2010; Yang et al., 2016). At the same time, research on model organisms has begun to illuminate the potential for brief stressors to have long-lasting effects, including on neuroendocrine function (Lupien et al., 2009) and the communities of internal microbes that are increasingly recognized as important components of organismal health (Bailey et al., 2011; Rea et al., 2016). These findings suggest that social interactions may change an individual’s phenotype in ways that alter their resistance to future stressors or recovery from past stressors. While these associations have been found in humans and some experimental work in lab-based model systems suggests the potential for causal relationships (Charuvastra & Cloitre, 2008; Holt-Lunstad et al., 2010; e.g., Yang et al., 2016), it is less clear how important the interaction between social conditions and the stress response is for wild animals living under real-world conditions.

A growing number of studies suggest that even under natural conditions, social environment might play an important role in mitigating or exacerbating responses to stressors, at least among mammals with complex social interaction networks (reviewed in Snyder-Mackler et al., 2020). Recent studies have demonstrated correlations in free living mammals between social experiences associated with dominance rank, social network integration, and affiliation or aggression and physiological measures associated with stress-coping, such as methylation patterns that alter glucocorticoid regulation (Snyder-Mackler et al., 2019), immune regulation (Snyder-Mackler et al., 2016), and gut microbiome composition (Tung et al., 2015). Even for species without complex social networks or dominance heirarchies, frequent social interactions mediated by signaling accumulate to result in biological differences that influence stress resilience. For example, in barn swallows (*Hirundo rustica*) the frequency and pattern of social interactions is correlated with microbiome diversity and corticosterone regulation (Levin et al., 2016). These physiological effects may in turn result in altered resilience to future stressors (Houtz et al., 2022; Taff & Vitousek, 2016).

Here, we simulated ecological challenges either before or after manipulating the experienced social environment in free-living tree swallows (*Tachycineta bicolor*). Challenges included either a series of simulated predation attempts or a wing surface reduction that produced an energetic challenge. Previous work on tree swallows has demonstrated that similar manipulations can result in mass loss, delayed breeding, and nest failure (Taff, Zimmer, et al., 2019; Winkler & Allen, 1995). Individual variation in resilience to these challenges is predicted by genome-wide DNA methylation (Taff, Campagna, et al., 2019) and a glucocorticoid stress response that includes a robust upregulation of corticosterone after capture coupled with strong negative feedback (Vitousek, Taff, Hallinger, et al., 2018; Zimmer et al., 2019).

Social environment was manipulated by dulling the white breast plumage that functions as a social signal. Naturally brighter white plumage in females from our study population is associated with the number and pattern of social interactions at the nest box and—under challenging conditions—naturally bright plumage is associated with higher reproductive success (Taff, Zimmer, et al., 2019). In other tree swallow populations, the brightness of white plumage is associated with immunity, nestling quality, nest retention, and the frequency or intensity of aggressive interactions at the nest box (Beck et al., 2015; Berzins & Dawson, 2016, 2018; Coady & Dawson, 2013). A previous experiment that manipulated color—but not ecological challenges—in this population found that experimental dulling changed the patterns of social interactions, increased the diversity of the cloacal microbiome, and resulted in higher overall reproductive success for dulled females, especially when those females were initially bright (Taff et al., 2021).

By manipulating both ecological challenges and the social environment, we sought to test the hypothesis that social conditions can alter the response to a realistic challenge. We predicted first that both ecological challenges and social manipulations would result in measurable differences in neuroendocrine function, the internal microbiome, and reproductive performance. Second, we predicted that the combination of social conditions and challenges would produce a different result than either treatment alone (e.g., that social conditions would result in priming that improved resilience to challenges). Finally, we tested whether the order of treatments (social manipulations before or after challenges) resulted in a different relationship between the two types of treatments.

## METHODS

### General Methods

We studied wild tree swallows breeding in nest boxes near Ithaca, New York, USA (42.503*°*N, 76.437*°*W) from May to July of 2018 and 2019. This population has been studied continuously since 1986 and standardized field methods for monitoring reproductive success are well established (Winkler et al., 2020). We conducted a separate experiment in each year, but general procedures for sampling birds and recording breeding activity were identical between the two years except where noted. Briefly, we monitored each nest box in the population every other day during the breeding season and every day around the expected hatching date to determine the timing of clutch initiation, onset of incubation (+/-1 day), and hatching (exact day).

Breeding females were captured three times each year: once around day 6 after the start of incubation, once near the expected hatching date (day 1 after hatching in 2018 and day 12 of incubation in 2019), and once about six days after hatching. We attempted to capture males six days after hatching, but since our experiments were focused on female-specific manipulations, we limited the length of male capture attempts to reduce overall disturbance; therefore, not all males were captured. Adults that were not banded in a previous year received a USGS aluminum band and a passive integrated transponder (PIT) tag encoding a unique ten-digit hexadecimal code for use with the RFID network (see below).

Female tree swallows have delayed plumage maturation and brown back plumage reliably indicates a ‘second year’ (one year old) female. Thus, for individuals that were not previously banded, we assigned age as second year (SY) or after second year (ASY) based on plumage. At the 1st and 3rd female captures and at male captures, we measured head + bill length (to the nearest 0.1 mm), wing length (to the nearest 0.5 mm), and mass (to the nearest 0.25 g). At the first capture, we collected 6-8 feathers from the center of the white breast to measure plumage brightness with a spectrophotometer as described in Taff et al. (2019). Treatments were applied at each capture as described for the two experiments below. At the 1st and 3rd capture in 2018, and at all three captures in 2019, we inserted a sterile flocked swab 1.5 cm into the cloaca (Puritan Medical Products Company LLC) to assay the cloacal microbiome following the procedure described in Taff et al. (2021). The swab was stored in 1 mL RNA later at −80° C until extraction.

Finally, we collected a series of three blood samples (1st and 3rd capture) or a single blood sample (2nd capture) to measure corticosterone. All blood samples were collected by brachial venipuncture into a heparinized micro-hematocrit tube. A baseline sample (< 70 *μ*l) was collected within 3 minutes of capture followed by a stress-induced sample (< 30 *μ*l) collected after 30 minutes of restraint. Immediately after the stress-induced sample was taken, we injected birds with 4.5 *μ*l/g of dexamethasone to stimulate negative feedback (Mylan 4mg ml*−*1 dexamethasone sodium phosphate, product no.: NDC 67457-422-00). Dexamethasone injection assesses the ability to downregulate corticosterone following a challenge and the procedure and dose have been previously validated in this population of tree swallows (Zimmer et al., 2019).

In 2019 (experiment two) at the 3rd capture only, rather than dexamethasone we injected females with Cortrosyn—a synthetic version of adrenocorticotropic hormone [ACTH]—to measure the ability of the adrenal cortex to respond to ACTH (50 *μ*l at 0.1 microgram per *μ*l; Amphastar Pharmaceutical Incorporated, Item # 054881). ACTH injection assesses the maximal physiological corticosterone response to a challenge. We validated this method using a separate set of individuals in a pilot experiment that is published in Taff et al. (2022). A final blood sample (< 30 *μ*l) was collected 30 minutes after injection. Immediately after blood sampling, glucose was measured in the field using a handheld FreeStyle Lite instant read glucose meter (Abbot Diabetes Care, Alameda, CA, USA; for validation data in tree swallows see Taff et al., 2021).

Our study was primarily focused on the behavior and physiological consequences of manipulations on adult female tree swallows. However, we also collected similar data and carried out an equal brood size cross fostering experiment on nestlings during these years. Primary analyses of nestling data are presented in a separate study (McNew et al., 2022). On day 12 after hatching, we banded nestlings with a USGS aluminum band, took the same morphological measurements as described above for adults, and took blood samples. The final fate of each nestling was determined by checking for fledging at 24 days after hatching. We used this fate to determine the number of nestlings fledged from each nest in the population.

All blood samples were stored on ice in the field for < 3 hours and then red blood cells and plasma were separated by centrifugation. Plasma was stored at −30°C until processing and red blood cells were stored in lysis buffer until DNA extraction. We measured corticosterone with enzyme immunoassay kits (DetectX Corticosterone, Arbor Assays: K014-H5) that were previously validated for tree swallows in this population (see supplementary methods for details on extractions and hormone measurements; Taff, Zimmer, & Vitousek, 2019).

### Experiment One: Signal Manipulation Followed by Challenge

In 2018, we carried out a 2×3 factorial experiment in which we first manipulated signal coloration, and then imposed an experimental challenge (see schematic of timeline for both experiments in Figure 1). At the first capture, females were alternately assigned a sham control or dulling treatment within each age group (ASY vs. SY). We chose to balance the treatment with age group because breeding phenology and reproductive success are known to differ between these age groups in tree swallows (Winkler et al., 2020). For this treatment, dulled females were colored across their entire white ventral surface with a light grey non-toxic marker (Faber-Castell PITT artist pen ‘big brush’ warm grey III 272). We previously validated that this treatment maintains the spectral characteristics of the plumage patch while reducing overall brightness (Taff et al., 2021). The treatment fades over time, but results in dulled plumage that lasts at least 10-14 days (Taff et al., 2021). Females in the sham treatment were colored for the same amount of time and over the same area using a clear marker (Prismacolor premier colorless blender PB-121). These treatments were re-applied at each of the subsequent two captures so that the plumage dulling persisted throughout the majority of the breeding season.

**Figure 1:**
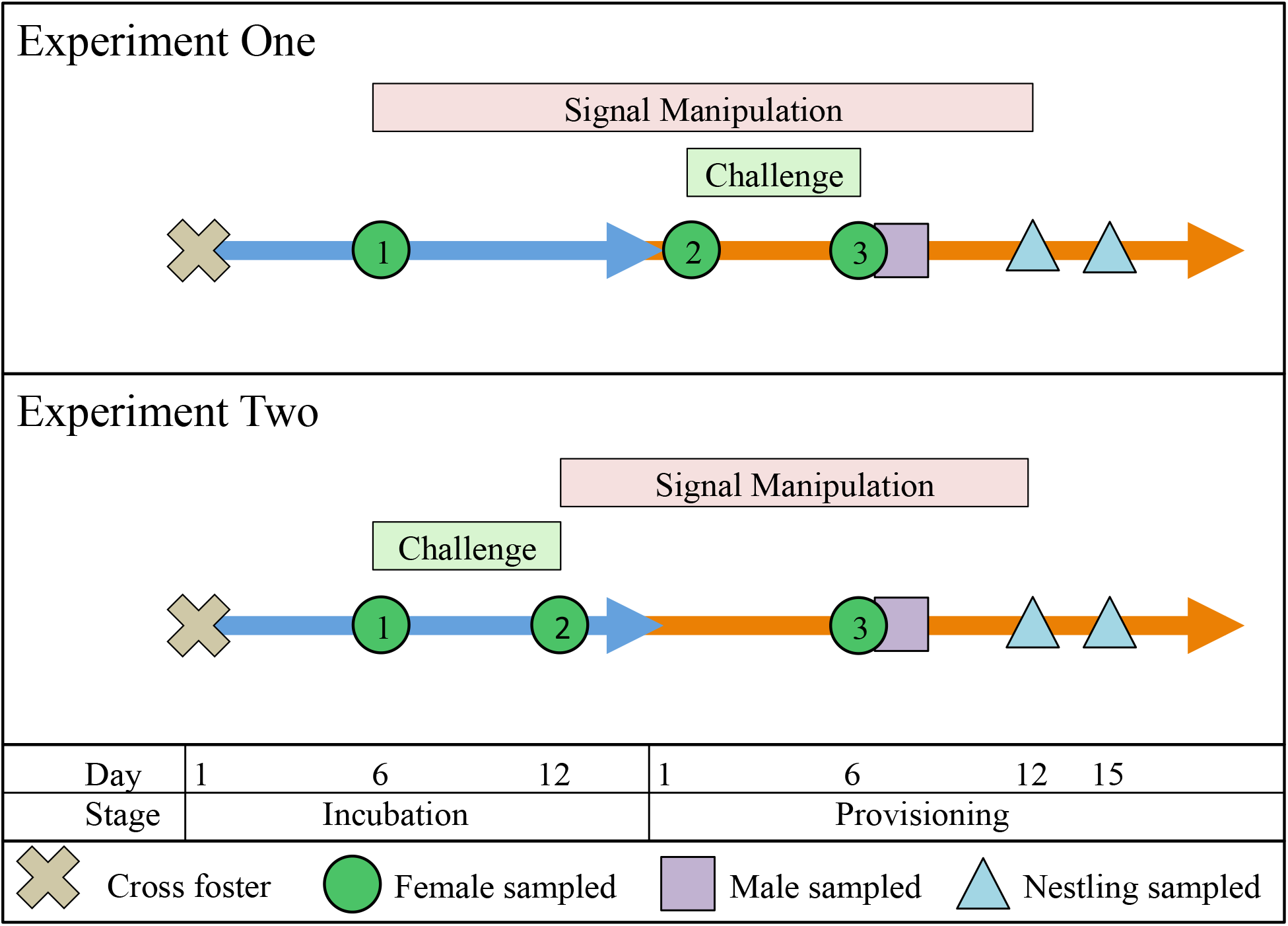
Schematic illustration of the experimental manipulations and measurements conducted in each year of study.

The second stage of this experiment involved imposing an experimental challenge that occurred between the 2nd and 3rd female captures. Within each signal manipulation group, females were alternately assigned a predation, flight handicapping, or control challenge treatment. As above, we attempted to balance each of these treatment combinations between age classes. For the predation treatment, we simulated two predation attempts on the female by a mink (*Neovison vison*), which is a common nest box predator on both adults and nestlings in the area of our field sites. Females were trapped in the nest box and then gently pulled out of the box using a taxidermied mink wrapped around a researcher’s hand. The bird was brought to the ground below the nest box and then allowed to escape. During these treatments, the researcher’s face and body were covered with a camouflage suit and the female was held facing away from the body to make the predation attempt seem as realistic as possible. This procedure was repeated two times during days 2-5 after hatching.

For the flight handicapping treatment, we used a modified version of a wing reduction treatment that was previously used to create an energetic challenge in this population (Taff, Zimmer, et al., 2019; Zimmer et al., 2019). Briefly, we bound together primary feathers 4, 5, and 6 on each wing with a miniature zip tie at the 2nd capture. This treatment reduces the wing surface area and therefore increases the energetic demands of flight and foraging (similar to the method used in Senar et al., 2002). The zip tie was removed at the 3rd capture, so that this energetic challenge lasted about 5 days. Finally, the control group received no additional treatment beyond the signal manipulation.

### Experiment Two: Challenge Followed by Signal Manipulation

In 2019, we carried out a 2×2 factorial experiment with the order of treatments from 2018 reversed. Due to a smaller number of available nests in 2019, we dropped the flight handicapping treatment from the 2019 experiment. At the first capture, we alternately assigned females to either a control or a predation treatment group within each age class. The predation group experienced simulated predation attempts identical to that described above for 2018, except that each female received three predation attempts and they occurred between days 8-12 of incubation.

All females were recaptured on day 12-13 of incubation and then alternately assigned to either a plumage dulling or sham signal manipulation treatment within each challenge treatment and age class. Signal manipulation proceeded exactly as described for 2018, with coloring re-applied at the third capture on day 6 after hatching.

### RFID Network for Provisioning and Social Behavior

We equipped every active nest box in our population with an RFID reader mounted under the box and connected to an antenna that circled the nest box entrance (described in Vitousek, Taff, Hallinger, et al., 2018). The reader was programmed to check for a PIT tag in the antenna every second and recorded activity from 5 am until 10 pm each day. Readers were installed four days after clutch completion and maintained throughout the breeding season. Some boxes, however, are missing RFID records in cases where a battery failed early or a reader malfunctioned. Therefore, the total number of days of RFID observation differs between boxes and we account for these differences in observation effort by modeling behavior on a daily basis using only days with full records (see details below).

We used data from the RFID sensor network to calculate both adult provisioning behavior and patterns of interaction between owners of different boxes in the population. From the raw RFID records, we determined provisioning behavior using a previously developed algorithm (Vitousek, Taff, Ardia, et al., 2018). Records were counted as unique provisioning trips if they occurred sufficiently far apart, with the time threshold depending on the age of nestlings based on a videotaped validation dataset from this population (Vitousek, Taff, Ardia, et al., 2018).

After identifying unique provisioning trips, we summarized provisioning effort into the number of unique trips per day for each adult. While we calculated provisioning data for both males and females, we primarily focus on female provisioning because treatments were targeted at females, because not all males were captured, and because males that were newly captured did not receive PIT tags until about 6 days after hatching. In this study, we included provisioning data from the day after hatching until nestlings were 15 days old. After day 15, parental provisioning declines quickly, nestling PIT tags can start to interfere with reading parental provisioning behavior, and the dulling treatment was expected to fade. Therefore, we considered day 15 the end of adult RFID data for all nests.

We also used raw RFID records to determine the number of times each day that focal birds were visited by other individuals in the population and the number of times that focal birds made trips to other boxes in the population. This procedure is described in detail in Taff et al. (2021). Because male and female visitation patterns differ (Taff et al., 2021) and because not all males had PIT tags for the entire breeding season, we focused only on visits to the focal box made by other females and on trips to other boxes made by the focal female. We considered visits or trips that occurred more than two minutes apart to be distinct records (see discussion in Taff et al., 2021).

### Microbiome Sample Processing and Bioinformatics

We processed swabs collected in the field to assess the cloacal microbiome exactly as described in Taff et al. (2021). Briefly, DNA was extracted with DNeasy PowerSoil kits (Qiagen Incorporated) following the manufacturer’s protocol and then the V4 region of the 16s rRNA gene was amplified in triplicate following the Earth Microbiome Project standard protocol (Caporaso et al., 2011, 2012). Samples were submitted to the Cornell Biotechnology Resources Center for sequencing in two Illumina MiSeq PE 2 × 250 bp runs (one run for each year). Raw sequences were processed following Callahan et al. (2016). At the end of this workflow, we calculated alpha diversity metrics for each sample using the `picante` and `phyloseq` packages in R with all samples rarefied to 1,000 reads (Kembel et al., 2010; McMurdie & Holmes, 2013). A full description of the lab work and bioinformatic pipeline is included in the supplemental methods.

### Data Analysis

As a general approach, we were interested in asking whether i) treatment had an effect on any of our behavioral, physiological, or performance outcomes and ii) whether the magnitude of a challenging experience was modified by an altered social interaction landscape (i.e., an interaction between the two treatment groups). Thus, our strategy was to fit models with the main effect of treatment group and an interaction between the two treatments as predictors. For some measures (e.g., 2nd capture physiology) the second half of the treatment had not yet been applied, and in these cases we fit models for the first stage of the experiment that included only fixed effects for the first half of the treatment sequence.

Based on previous work in this population, we expected initial breast brightness to modulate the effect of the dulling treatment on social activity because initially bright birds will experience a greater reduction in brightness when colored (Taff et al., 2021). We also expected initial brightness to be associated with reproductive success (Taff, Zimmer, et al., 2019). Therefore, we included an interaction between initial coloration and dulling treatment in models of provisioing, social interaction, and reproductive success, but we removed this effect when there was no support for initial brightness to simplify interpretation. While initial brightness might also contribute to differences in other performance measures, our sample size was insufficient to reliably model three-way interactions between initial brightness and both treatment levels, so we primarily focused on simpler models built around the main effects of the two treatments. The exact form of our models differed depending on the response variable in question. For adult morphology, glucose, corticosterone, and microbiome diversity analyses, we fit linear mixed models that included an interaction between the two treatments and the stage (incubation or nestling, see below) along with the initial, pre-treatment measure for the response variable in question when available.

Data extracted from the RFID network on provisioning and visitation patterns included multiple days of observation for each nest. For provisioning data, we fit linear mixed models with the total number of provisioning trips made by the focal female each day as the response variable. Predictors included the two treatments and their interaction with pre-treatment female brightness (dropped if not supported) along with nestling age and brood size (as in Vitousek, Taff, Ardia, et al., 2018). Because provisioning increases with nestling age, we also included an interaction between nestling age and treatment to allow for a different slope of increase in each treatment. Random effects were included for female identity to account for repeated observations and for the day of year to capture variation in provisioning associated with weather or overall food availability not related to our treatments (see Vitousek, Taff, Ardia, et al., 2018 for further discussion).

For social interactions, we focused on the number of visits to the focal nest box by other females in the population and trips made to other nest boxes in the population on each day of observation. For each of these two metrics, we fit separate models for stage one (mainly during incubation) and stage two (mainly during nestling provisioning) periods. We split the stages in our models because they differ both in which stage of treatment had been applied already and—importantly—in the overall frequency of social interactions; visits to other nest boxes are much more common during the nestling provisioning stage (Taff et al., 2021). Each model included the one (stage one) or two (stage two) treatments that had been applied along with initial brightness (dropped if not supported) as fixed effects. Models also included the day in the breeding cycle (0 = hatching day) as a linear predictor. Random effects included individual identity and day of year as above for provisioning. These social interaction models were fit as generalized linear mixed models with a Poisson distribution.

Finally, we fit generalized linear mixed models with a binomial distribution to look for differences in fledging success for each treatment group (modeled using the fate of each individual nestling). These models included the two treatment stages and their interaction along with initial female brightness. Brightness was dropped if the effect was not supported. The models also included a random effect for nest identity. We assessed the support for either interactions between the treatments or main effects of each treatment by comparing marginal effects using the `emmeans` package where appropriate (Lenth, 2020). Note that more detailed analyses of the effect of treatments and cross fostering on nestling development and physiology are included in separate studies (McNew et al., 2022; Taff, Johnson, et al., 2022).

All analyses and figures were produced in R version 4.0.2 (R Core Team, 2020). Mixed models were fit using the ‘lmer’ or ‘glmer’ functions in the R package ‘lme4’ (Bates et al., 2014). We interpreted effects with 95% confidence intervals that did not cross zero as meaningful; p-values based on the Satterthwaite method as implemented by ‘lmerTest’ in R are reported in the supplementary materials (Kuznetsova et al., 2017). Any continuous predictor variables were scaled to mean of 0 and standard deviation of 1 to make comparison of coefficients easier. We ran each set of models separately for the two independent experiments and report results from each experiment side by side for comparison. Full details on each fit model are included in the supplementary materials. A complete set of annotated scripts and data required to reproduce all analyses and figures presented in the article and in the supplemental material is permanently archived at Zenodo (DOI: 10.5281/zenodo.7506939). Raw microbiome sequence data are publicly available through the NCBI Short Read Archive (PRJNA918663).

## RESULTS

In experiment one (2018) we sampled a total of 57 female breeding attempts (sample size by treatment group: sham + control = 10; sham + predator = 11; sham + handicap = 9; dulled + control = 8; dulled + predator = 10; dulled + handicap = 9). In experiment two (2019) we sampled a total of 54 breeding attempts (sample size by treatment group: control + sham = 15; control + dulled = 15; predator + sham = 12; predator + dulled = 12). The exact sample sizes used in each comparison below differed depending on data availability for each measurement.

### Daily Social Interaction Rates

In experiment one, the first stage of treatments, in which signals were manipulated, occurred from day six of incubation to day one after hatching. During this stage, there was no difference between experimentally dulled and sham colored females in the number of visitors received at the nest box or in the number of trips that experimental birds made to other boxes (Table S1). The second stage of treatments, in which experimental challenges were applied, lasted from day one to fifteen after hatching; there was still no difference in the number of trips made to other boxes during this stage (Table S1). However, during this stage the number of visits that experimental nests received from other females depended on both treatment and initial brightness of the focal female (Figure 2; Table S1). Females in the control group (where neither plumage brightness nor challenge exposure was manipulated) that had bright breast plumage received more daily visits by other females to their nest than naturally darker control females did (Figure 2). This same pattern was observed in the group that received signal dulling plus simulated predation or handicap treatments, although in the dulling plus handicap group the confidence interval for the slope overlapped zero (Figure 2). However, there was no relationship between initial brightness and visit rate for females that received simulated predation or handicapping with no signal dulling or for females that received signal dulling with no challenge (Figure 2).

**Figure 2:**
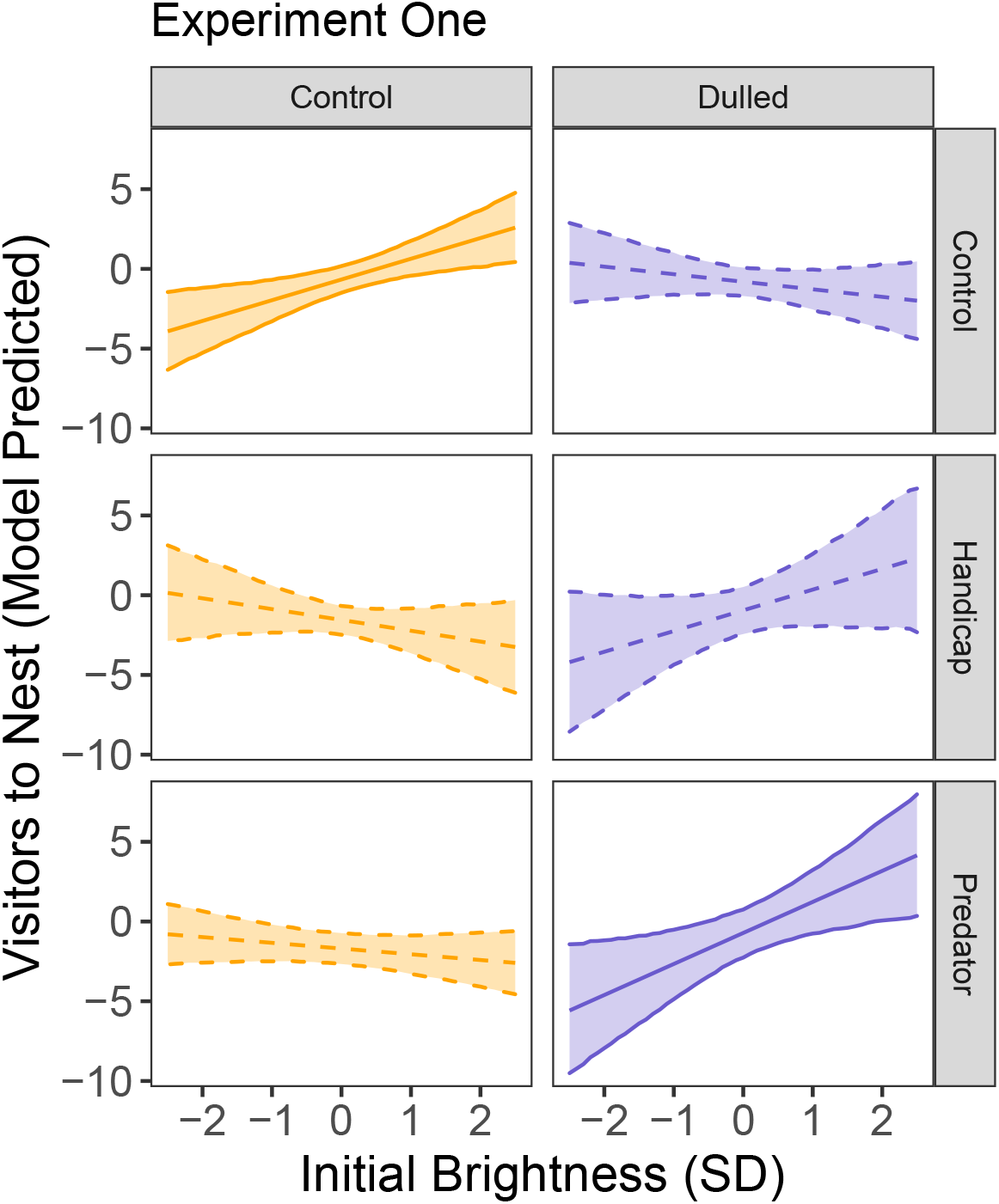
Model predicted relationship between initial brightness and the number of daily visits to the nest by other females for each treatment group during the second stage of experiment one. Central lines are the maximum likelihood estimates and shaded regions are 95% confidence intervals. Solid lines indicate relationships in which the 95% CI of the slope did not span zero, whereas dashed lines indicate relationships where the slope estimate crossed zero.

In experiment two, the first stage of the treatment was simulated predation and it occurred from day six to twelve of incubation. There was no evidence that the number of visitors to the box or trips to other boxes differed by treatment group in this period (Table S2). The second stage of treatments occurred from day twelve of incubation to day fifteen after hatching. During this period, the number of visitors to the box was still unrelated to treatments (Table S2). However, in the signal dulling plus predator treatment only there was a negative relationship between initial brightness and the number of daily trips made to other boxes (Figure 3).

**Figure 3:**
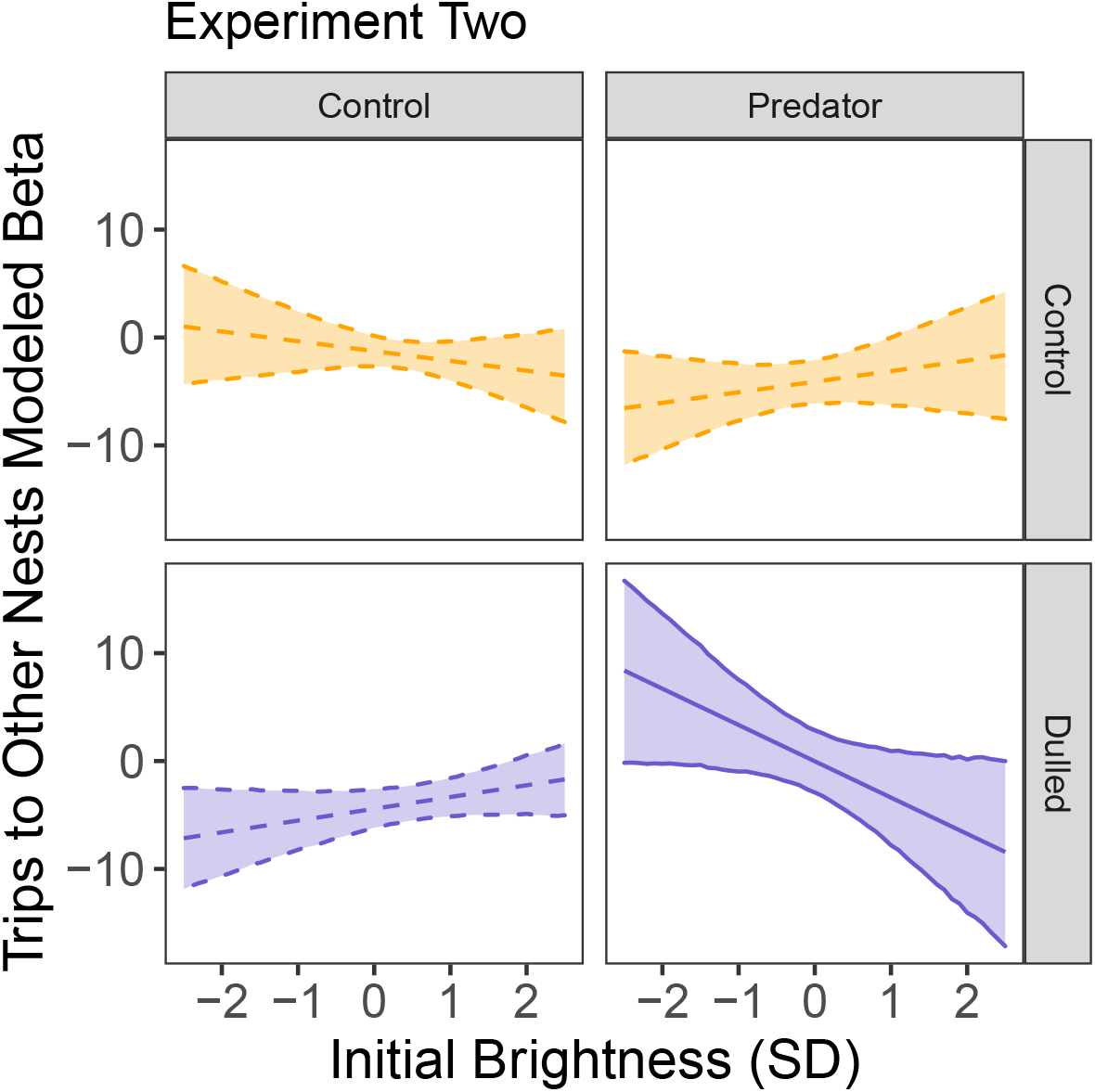
Model predicted relationship between initial brightness and the number of daily trips to other boxes by the focal female during the second stage of experiment two. Central lines are the maximum likelihood estimates and shaded regions are 95% confidence intervals. Solid lines indicate relationships in which the 95% CI of the slope did not span zero, whereas dashed lines indicate relationships where the slope estimate crossed zero.

### Female Mass and Physiology

In both years, female mass declined across the three captures and pre-treatment mass predicted mass later in the season (Table S3; experiment one effect of breeding stage *β* = −1.53, 95% CI = −2.49 to −0.57, effect of initial mass *β* = 0.6, CI = 0.36 to 0.85; experiment two effect of breeding stage *β* = −2.02, CI = −2.72 to −1.32, effect of initial mass *β* = 0.7, CI = 0.46 to 0.95). However, there was no evidence that any treatment or combination of treatments altered this seasonal decline in mass in either year (Table S3).

Although there were some correlations between different corticosterone measurements within females, no corticosterone measurements were related to treatment in either year (Table S4). Similarly, baseline and stress induced glucose measurements were unrelated to treatments in either year (Table S5).

### Female Microbiota

Overall, the most common phyla detected in the cloacal microbiome were Actinobacteria (39 % of total amplicon reads), Proteobacteria (33.1 %), Firmicutes (9.9 %), and Tenericutes (9.6 % Figures S1 & S2). In experiment one, there was some evidence that alpha diversity differed by treatment groups, but these effects were weak and sensitive to the choice of alpha metric. At the 3rd capture, females in the flight reduction treatment had higher Shannon diversity (Table S6; effect of flight reduction *β* = 0.82, CI = 0.07 to 1.57). At this capture, treatment was also related to Inverse Simpson diversity; females in the signal dulling treatment had higher Inverse Simpson diversity, but this relationship was strongest among dulled females that did not also receive the predation treatment (Table S6; main effect of signal dulling *β* = 4.89, CI = 0.85 to 8.94, interaction between dulling and predator treatment *β* = −5.33, CI = -11.16 to 0.51). In experiment two, there was no evidence that treatment group was related to alpha diversity at either the 2nd or 3rd capture, regardless of the metric used (Table S6).

Visual inspection of ordination plots showed little evidence for separation of microbial communities between treatment groups in either experiment one (Figure S3) or experiment two (Figure S4). We tested for differences in the centroid or dispersion of communities using the ‘adonis2’ function from package vegan in R using the default settings (Oksanen et al., 2013). In experiment one, there was no difference between groups (all effects P > 0.2). In experiment two, there was no difference between groups at the 1st or 2nd capture (all effects P > 0.25), but at the third capture there was a significant interaction between signal dulling and predator treatments (PERMANOVA: dulling F_1,35_ = 1.54, P = 0.10; predator F_1,35_ = 0.66, P = 0.84; dulling*predator F_1,35_ = 2.13, P = 0.03). A subsequent test of differential dispersion using the ‘betadisper’ function indicated that this difference was likely driven by a difference in dispersion between the predator treatment groups that did or did not receive signal dulling (betadisper effect of treatments F_3,35_ = 3.26, P = 0.04). Inspecting the ordination plot (Figure S4) suggests that the small within group dispersion in the sham plus predator treatment group drove this difference rather than any difference in the multivariate centroid of microbial communities between treatments.

### Daily Provisioning Rates

In experiment one, the total daily number of female provisioning trips increased as nestlings aged (main effect of nestling age *β* = 7.09, CI = 4.80 to 9.37). There was some support for an interaction between nestling age and treatment and this was mainly driven by a shallower increase in provisioning rate as nestlings aged in the sham color compared to the sham color plus predator treatment group (Figure 4; Table S7).

**Figure 4:**
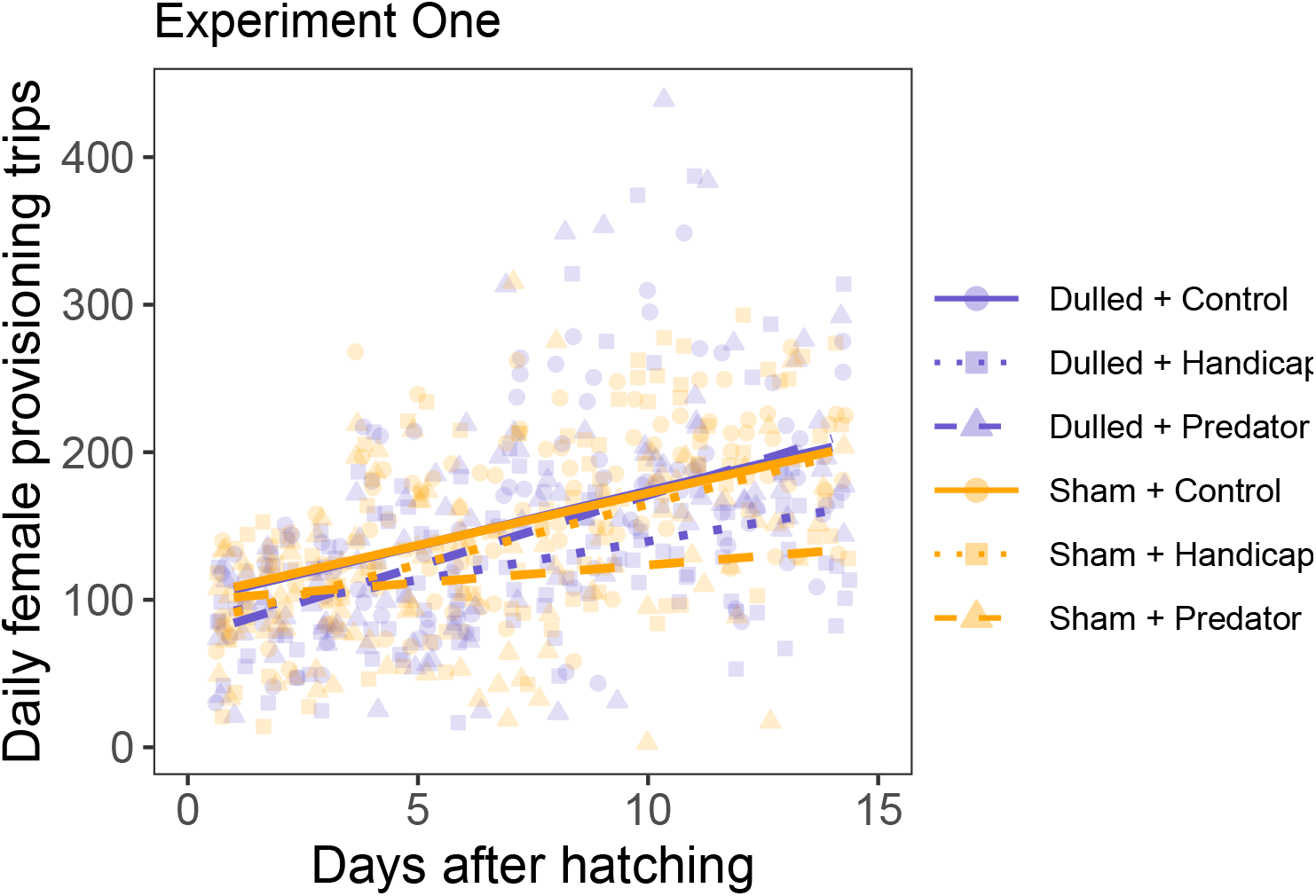
Total female provisioning trips by nestling age and treatment in experiment one. Lines show the model predicted change in provisioning with nestling age for each treatment. Faded points are the raw data points for each individual female on each day. See text for description of the model parameters.

In experiment two, daily female provisioning showed a similar relationship with nestling age and was also higher in nests with larger broods (Table S7; main effect of brood size *β* = 20.88, CI = 12.92 to 28.85; nestling age *β* = 6.11, CI = 3.39 to 8.82). Females in the plumage dulling treatment tended to increase provisioning with nestling age at a steeper rate, but the confidence intervals for each group overlapped (Figure 5; Table S7). Predator treatment was not related to provisioning rate (Figure 5).

**Figure 5:**
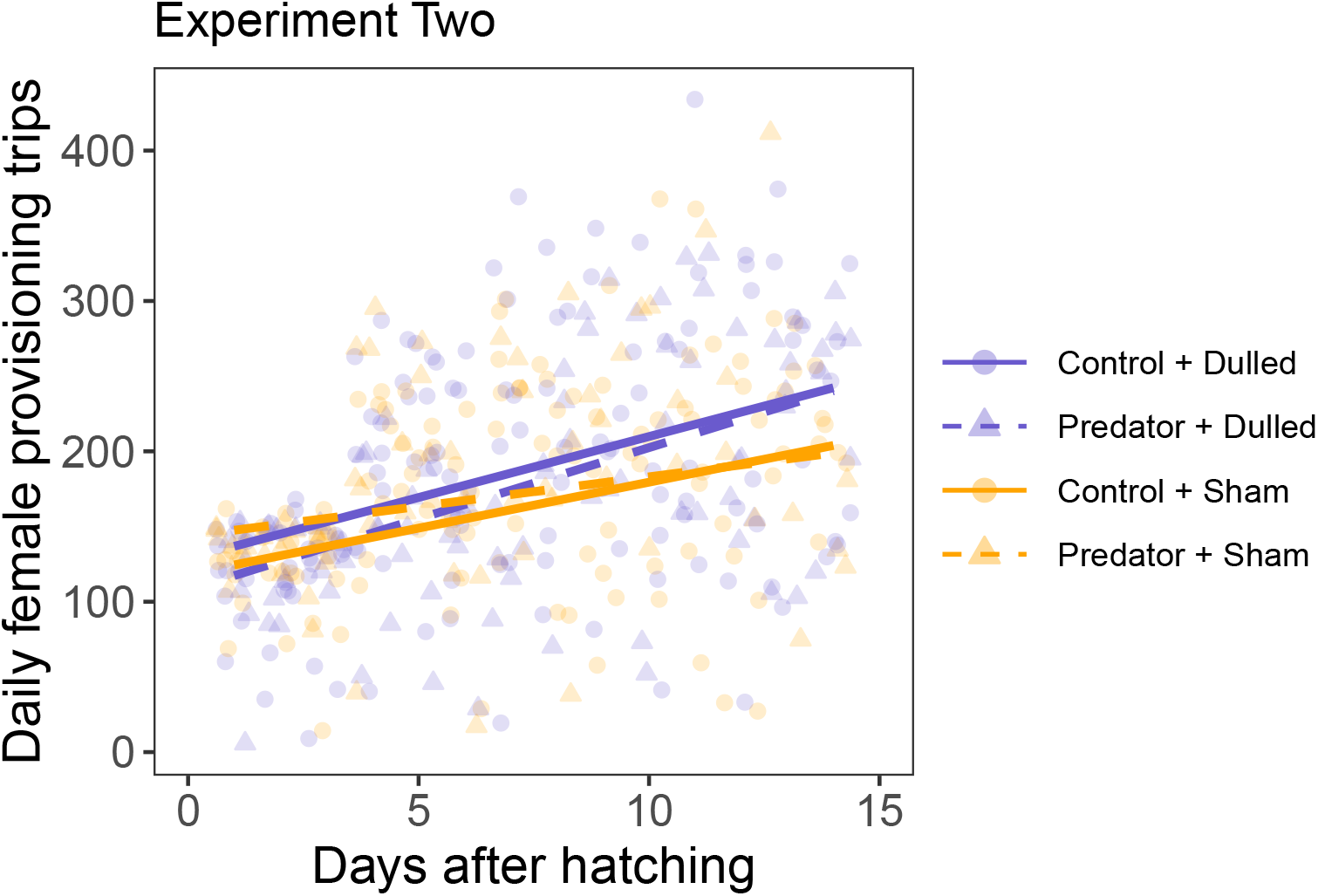
Total female provisioning trips by nestling age and treatment in experiment two. Lines show the model predicted change in provisioning with nestling age for each treatment. Faded points are the raw data points for each individual female on each day. See text for description of the model parameters.

### Fledging Success

In experiment one, the likelihood of fledging was lower for nestlings in the predator treatment (Table 1 & S8; `emmeans` marginal effect for control *β* = 0.55, CI = −0.43 to 1.53; predator *β* = −1.78, CI = −2.80 to −0.75; handicap *β* = −0.34, CI = −1.28 to 0.60). However, there was no difference between sham and signal dulling groups and no support for an interaction between dulling and challenge treatment (Table S8). In experiment two, there was no difference in the likelihood of fledging between any treatment groups, although fledging likelihood was lower in nests at which females were initially bright (Table 1 & S8).

**Table 1:** Average number of nestlings fledged by treatment group.

| First Treatment | Second Tretament | Year | Sample Size | Number Fledged | SD |
| --- | --- | --- | --- | --- | --- |
| Sham | Control | 2018 | 11 | 3.0 | 2.2 |
| Sham | Predator | 2018 | 11 | 1.4 | 1.7 |
| Sham | Handicap | 2018 | 11 | 2.8 | 1.9 |
| Dulled | Control | 2018 | 9 | 3.2 | 1.6 |
| Dulled | Predator | 2018 | 11 | 1.3 | 1.5 |
| Dulled | Handicap | 2018 | 10 | 2.2 | 2.0 |
| Control | Sham | 2019 | 17 | 2.6 | 2.1 |
| Control | Dulled | 2019 | 16 | 1.9 | 2.2 |
| Predator | Sham | 2019 | 14 | 1.0 | 2.0 |
| Predator | Dulled | 2019 | 15 | 1.7 | 1.8 |

## DISCUSSION

In this study, we found that experimental challenges and dulling of a social signal impacted fledging success and social interactions, respectively. While there was some evidence that both experimental challenges and signal manipulations influenced the social environment that females experienced, there was no clear evidence for an interaction between the two categories of manipulation on physiological measurements or fledging success. Females that were subjected to simulated predation events after their eggs hatched had nestlings with lower fledging success, but the effect did not depend on signal dulling. Females with experimentally dulled signals or facing experimental challenges experienced an altered social environment, but this had no apparent effect on resilience to subsequent challenges. Likewise, there was no evidence that the order of manipulations altered their effects, though for both categories of manipulation the effects appeared to differ when applied during incubation compared to the nestling period. Taken together, our results do not support our main hypothesis that alterations to the social environment modulate resilience to a subsequent challenge, at least for this combination of treatments and under the environmental conditions that we observed. However, as discussed below, the overall effect of challenge treatments was weaker than expected. Thus, it is possible that the effect of the social environment would differ for individuals coping with significant challenges.

The effects of our predator manipulations broadly align with previous work suggesting that non-consumptive effects of predation can result in differences in seasonal reproductive success (Clinchy et al., 2013; e.g., Lima, 2009; Zanette et al., 2011). While several studies in other species have found that predator cues or encounters can alter parental provisioning rates (LaManna & Martin, 2016) and nestling mass or physiology (Ibáñez-Álamo et al., 2011), we found no evidence here that provisioning rates or nestling measurements differed in predator-treated nests. It is possible that mortality biased towards the smallest nestlings in predator-treated nests could have obscured these effects, but we could not test for this possibility since nestlings were not measured until day 12 after hatching when some had already died. Thus, it is unclear exactly why fewer nestlings fledged from predator-treated nests, though it seems likely that mortality soon after hatching played a role. We only found a clear effect of predator treatments on fledging success when the treatments occurred during the nestling period, suggesting that the effect was sensitive to timing in the breeding cycle (see McNew et al., 2022 for further discussion). Although it is worth noting that even in experiment two, nests that received the predation treatment only fledged an average of 60% of the nestlings as control nests, but given the variation and sample size this difference was not statistically distinguishable.

In several small songbirds, predator cues during incubation can result in altered incubation or provisioning behavior that might impact nestling development and survival (LaManna & Martin, 2016). We saw no clear evidence that provisioning rate differed with predator treatment, but one possibility that we could not assess with our data is that females may have spent less time brooding after experiencing simulated predation attempts. Thermal environment during development is known to be important for tree swallows (Shipley et al., 2020, 2022; Uehling et al., 2020) and could have resulted in different overall energy expenditures for nestlings in predator-treated nests if nestlings were more exposed to ambient temperature because of reduced parental brooding. While some studies have suggested that nestlings might directly respond to predator cues (Ibáñez-Álamo et al., 2011; Zwaan & Martin, 2020), there was no evidence for these effects in our study (McNew et al., 2022).

Surprisingly, we saw little effect of the flight efficiency reduction treatment employed in experiment one. A similar treatment was previously found to result in increased nest failure, but females with strong negative feedback and bright plumage were more resilient to the challenge (Taff, Zimmer, et al., 2019; Zimmer et al., 2019). This study had two important differences that might explain the discrepancy in results. First, the previous studies were conducted during a breeding season that had an unusually harsh cold snap that resulted in extensive nestling mortality (Shipley et al., 2020), whereas the years of study for these experiments were relatively benign. Cold snaps reduce food availability for tree swallows (Winkler et al., 2013) and often result in nestling mortality (Shipley et al., 2020); thus the combination of a cold snap and flight handicapping might be an especially severe challenge. Second, in an effort to moderate the challenge imposed by the treatment, we reduced the surface area reduction of feathers in this experiment compared to our prior study. The combination of these two differences may have resulted in a treatment that was not sufficiently challenging to influence breeding performance. That being said, there was some evidence that microbiome alpha diversity differed for birds in the flight reduction group, although this result depended on the metric chosen. The marginal change in microbiome alpha diversity that we detected might suggest that flight handicapping did have some effect on diet or energy budget, albeit a small one.

The social consequences of signal dulling in experiment one were somewhat similar to those observed in a previous study of this population that employed the same dulling technique without subsequent challenges (Taff et al., 2021), although in the present study the effects were less pronounced. In experiment two, there was less clear evidence for an effect on social behavior, although females in the dulling plus predator group made fewer trips to other boxes if they were initially bright. Despite the fact that these social interaction patterns were supported statistically, none were particularly clear or strong and all depended on interactions with initial brightness. It is unclear why our results differed from previous observational (Taff, Zimmer, et al., 2019) and manipulative (Taff et al., 2021) studies, but there are a few possible explanations. First, we know that overall social interaction rates differ substantially from year to year, probably in relation to local weather conditions and breeding density that directly impact breeding behavior (Shipley et al., 2022; Taff et al., 2021). Compared to prior studies, the two years that these experiments were conducted in had relatively benign weather but relatively low breeding density. Second, the timing of our dulling treatments differed somewhat from our previous study, particularly in experiment two in which the dulling was not applied until late incubation. It is possible that the effect of changing color later in the breeding season was smaller because there had already been more time to directly assess neighbors. In any case, the fact that the effects on social behavior were less pronounced than expected means that we cannot assess whether a more successful social manipulation would have resulted in an interaction with challenge treatments.

Although dulling had some effect on social interactions, there was relatively little evidence for downstream consequences of social interactions or dulling on reproductive performance. Our previous study found that dulled females provisioned nestlings at a higher rate, had a more diverse microbiome, and fledged more offspring (Taff et al., 2021). Here, we again found that dulled females provisioned more, but only in the second experiment and the effect was only marginally supported. Similarly, we found that dulled and flight handicapped females had higher microbiome alpha diversity, but only in first experiment. Moreover, this effect was weak and sensitive to the exact alpha diversity metric used. In contrast, there was no evidence that signal dulling was related to any other measures of physiology or to ultimate fledging success. Because the effect of our social signal manipulation was less dramatic than in our previous study, at this time we cannot say whether these differences in downstream effects are the consequence of a less intense social manipulation or whether social interactions are not causally linked to the downstream outcomes that we measured.

While there is growing interest in understanding the effects of social interactions on resilience to challenges, relatively few studies have addressed these questions in natural populations (but see, Anderson et al., 2021, 2022; Snyder-Mackler et al., 2019). At present, it is unclear whether results from lab rodents with simplified social environments or from primates that have highly developed social interaction systems will be generalizable to other species. It is possible that in many cases, the subtle effects of social interactions on stress resilience may be swamped by natural variation in abiotic conditions that have larger magnitude effects on coping ability. The variability that we observed in the effect of similar social signal manipulation between years in tree swallows suggests that this may have been the case in our study.

Despite the fact that our study did not find any clear evidence for an interaction between the social environment and ecological challenges, we suggest that it is too early to say how important these interactions might be in many species, including in tree swallows. In the case of our study, the lack of interaction might have resulted from either the relatively weak effect of each treatment alone, the exact choice and timing of treatments we used, or a true lack of interaction. Distinguishing between these possibilities will be challenging and in many cases species-specific biology may be important. Nevertheless, we suggest that empirical work in this area could benefit from the continued development of conceptual and mechanistic models that explicitly consider the conditions under which the social environment is predicted to affect resilience, the characteristics of challenges that may trigger environment-dependent responses, and what downstream consequences of challenges are most likely to show biological evidence of these interactions.

## ETHICAL NOTES

All procedures were approved by the Cornell University Institutional Animal Care & Use Board (IACUC protocol # 2019-0023). All work was conducted with appropriate state and federal permits to MNV.

## ACKNOWLEDGMENTS

We thank the field and lab technicians who helped collect data for this project, including Bashir Ali, Allison Anker, Paige Becker, Raquel Castromonte, Jeremy Collison, KaiXin Chen, Alex Dopkin, Zapporah Ellis, Audrey Fox, Brianna Johnson, Christine Kallenberg, Raisa Kochmaruk, Alex Lee-Papastravos, Jabril Mohammed, Yusol Park, Alyssa Rodriguez, Bella Somoza, and Kwame Tannis. We also thank Brittany Laslow and Benj Sterrett for support in the lab and field.

## SUPPLEMENTARY MATERIAL FOR

### SUPPLEMENTAL METHODS

#### Plumage Measurement

We measured the overall brightness of feathers collected from the center of the white breast of each female in this study exactly as described in previous work on this population (Taff, Zimmer, et al., 2019; Taff et al., 2021). Briefly, four feathers from the breast were stacked and taped onto black construction paper. Reflectance was then measured with an Ocean Optics FLAME-S-UV-VIS spectrophotometer with PX-2 pulsed Xenon light source and WS-1 white standard in OceanView version 1.5.2 (Ocean Optics, Dunedin, FL). Aquisition settings included a 20 nm boxcar width, 10 scan average, and 60 ms integration time. We used a holster on the fiber optic UV/VIS probe so that light was blocked while reading and the probe was held a constant 5 mm distance from the feathers. For each feather stack, we took four separate spectral readings with the probe removed between each reading.

We processed raw reflectance spectra using the `pavo` package in R (Maia et al., 2013). As in Taff et al. (2021), we focused on the overall brightness of the breast plumage given by the ‘B2’ measurement in `pavo`. This measurement represents the average reflectance across the range of 300-700 nm. Finally, the brightness measurements from the four separate spectra were averaged for each individual to arrive at a single brightness measure for each individual in the population before treatments were applied. In this study, we did not collect additional feathers at later time points after treatments, but Taff et al. (2021) includes extensive validation data with feathers measured at varying time points after treatments for sham and dulled birds.

#### Corticosterone Measurement

We measured the concentration of corticosterone from plasma that was frozen in the field using commercially available enzyme immunoassay (EIA) kits (DetectX Corticosterone, Arbor Assays: K014-H5). Validation and lab testing data on these kits when applied to our population of tree swallows is available in Taff et al. (2019). We first extracted corticosterone from plasma by adding 5 *μ*l of plasma to 45 *μ*l of assay buffer and then proceeding with three rounds of ethyl acetate extraction. The final extract was dried overnight in a fume hood and then reconstituted with 125 *μ*l of assay buffer. Reconstituted samples were run in duplicate with a 9 point standard curve in 96 well EIA plates. Average extraction efficiency was determined by spiking some samples with a known amount of concentrated corticosterone and determining the percent recovery.

Using these starting volumes, the lower detection limit for corticosterone was 0.8 ng/*μ*l and we substitute this value for any samples that were too low to detect. Overall, the extraction efficiency was 96.2%. When comparing replicates of each sample run on the same plate, the intra-plate CV was 11.5%. The inter-plate CV, based on plasma pools run across multiple plates, was 12.7%.

#### Microbiome Lab Procedures

Samples were collected in the field with flocked sterile swabs (Puritan® HydraFlock®; product number 25-3317-H) that were inserted ∼1.5 cm into the cloaca and then gently removed while slowly rotating (as in Vo & Jedlicka, 2014). We stored samples in 1 mL of RNAlater at −80 C and then extracted DNA according to the manufacturer’s protocol with DNeasy PowerSoil DNA Isolation Kits (Qiagen Incorporated). After extraction, we used the 515F and 806R primers with Illumina adapters to amplify the V4 region of the 16S gene following the standard Earth Microbiome Project protocol modified for 10 *μ*l reactions (Caporaso et al., 2011, 2012). In experiment one, we collected microbiome samples only at the first and third captures, while in the second experiment we collected samples from all three captures.

We ran each PCR reaction in triplicate and succesful amplification was confirmed by running each pooled reaction on a 1% agarose gel. Pooled samples were submitted to the Cornell Biotechnology Resource Center for quantifiaction, library preparation, and sequencing (all lab procedures as described in Taff et al., 2021). Each sequencing run also included negative control extractions that went through the full procedure from extraction to sequencing with only a sterile swab and RNAlater. Extracted samples from adults were sequenced in two lanes (one each for the 2018 experiment and the 2019 experiment) on an Illumina MiSeq PE 2 × 250 run.

#### Microbiome Bioinformatic Pipeline

We processed sequences with the exact same workflow described in Taff et al. (2021), which was based on a published workflow for processing microbiome sequences by Callahan et al. (2016). All forward and reverse reads were truncated to 180 bp using the ‘filterAndTrim’ function in the `dada2` package in R (Callahan, McMurdie, et al., 2016). After trimming, we proceeded through dereplication, modeling sequencing errors, determining amplified sequence variants (ASVs), merging reads, and removing chimeras all using the default settings in `dada2` in order to create a final set of ASVs for each sample.

We used the Silva 132 database to make taxonomic assignments (Quast et al., 2012; Yilmaz et al., 2014) and built a generalized time-reversible maximum likelihood tree from the ASVs using the `phangorn` package in R (Schliep, 2011). Subsequent analyses were carried out using the `phyloseq` and `vegan` package in R to combine the ASV table with sample data, ASV taxonomy, and the phylogenetic tree (Dixon, 2003; McMurdie & Holmes, 2013). After removing singletons and non-bacterial ASVs, we processed reads with the `decontam` package to identify and remove likely contaminants (Davis et al., 2018).

We used the final ASV file to determine the relative abundance of taxa in each sample. For our main analysis, we calculated alpha diversity metrics for each sample after agglomerating taxa at the genus level (Shannon Index, Faith’s Phylogenetic Diversity [PD], and Inverse Simpson’s Index) using the `phyloseq` and `picante` packages in R (Kembel et al., 2010; McMurdie & Holmes, 2013). When calculating alpha diversity metrics, we rarefied all samples to 1000 reads using the ‘prune_samples’ function in `phyloseq`.

We tested for differences in alpha diversity across treatment groups using linear models that included alpha diversity at the second or third capture as the response along with first capture diversity and an interaction between the two treatment levels as predictors. In addition to comparing alpha diversity metrics, we also tested for dissimilarity in the microbiota community composition between the treatment groups using the `vegan` package in R (Dixon, 2003). For these analyses, we calculated Bray-Curtis dissimilarity matrices using principal coordinate analyses (PCoA). Differences between treatment groups were assessed by PERMANOVA tests as implemented by the `adonis2` function in `vegan`. When significant effects were detected with PERMANOVA, we moved on to dispersion tests with the `betadisper` function to determine whether differences were driven by within-group dispersion.

## SUPPLEMENTAL FIGURES AND TABLES

**Figure S1:**
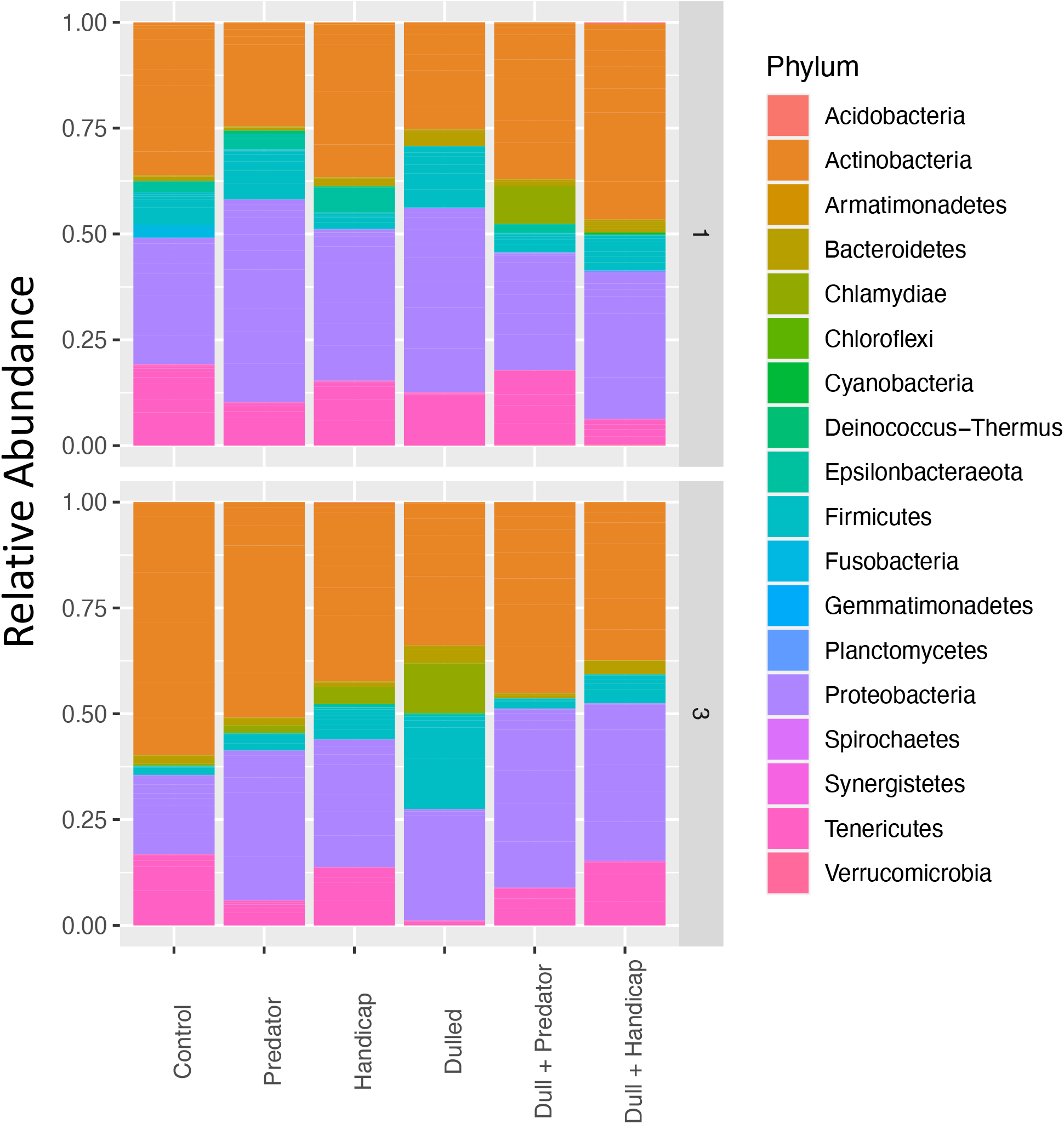
Relative abundance based on number of reads for bacterial phyla in samples from the first (pre-treatment) and third (post-treatment) capture in experiment one.

**Figure S2:**
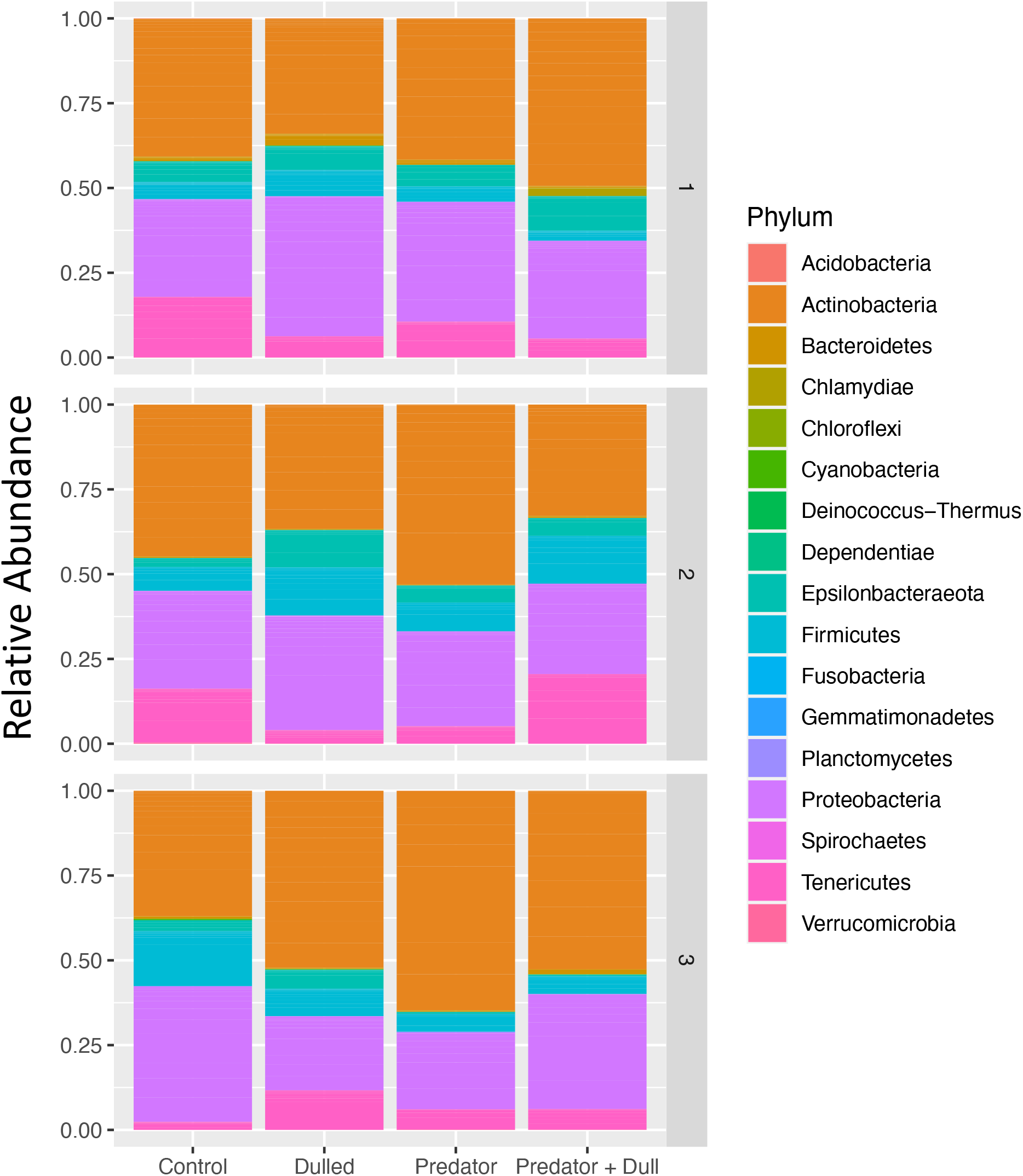
Relative abundance based on number of reads for bacterial phyla in samples from the first (pre-treatment), second, and third (post-treatment) captures in experiment two.

**Figure S3:**
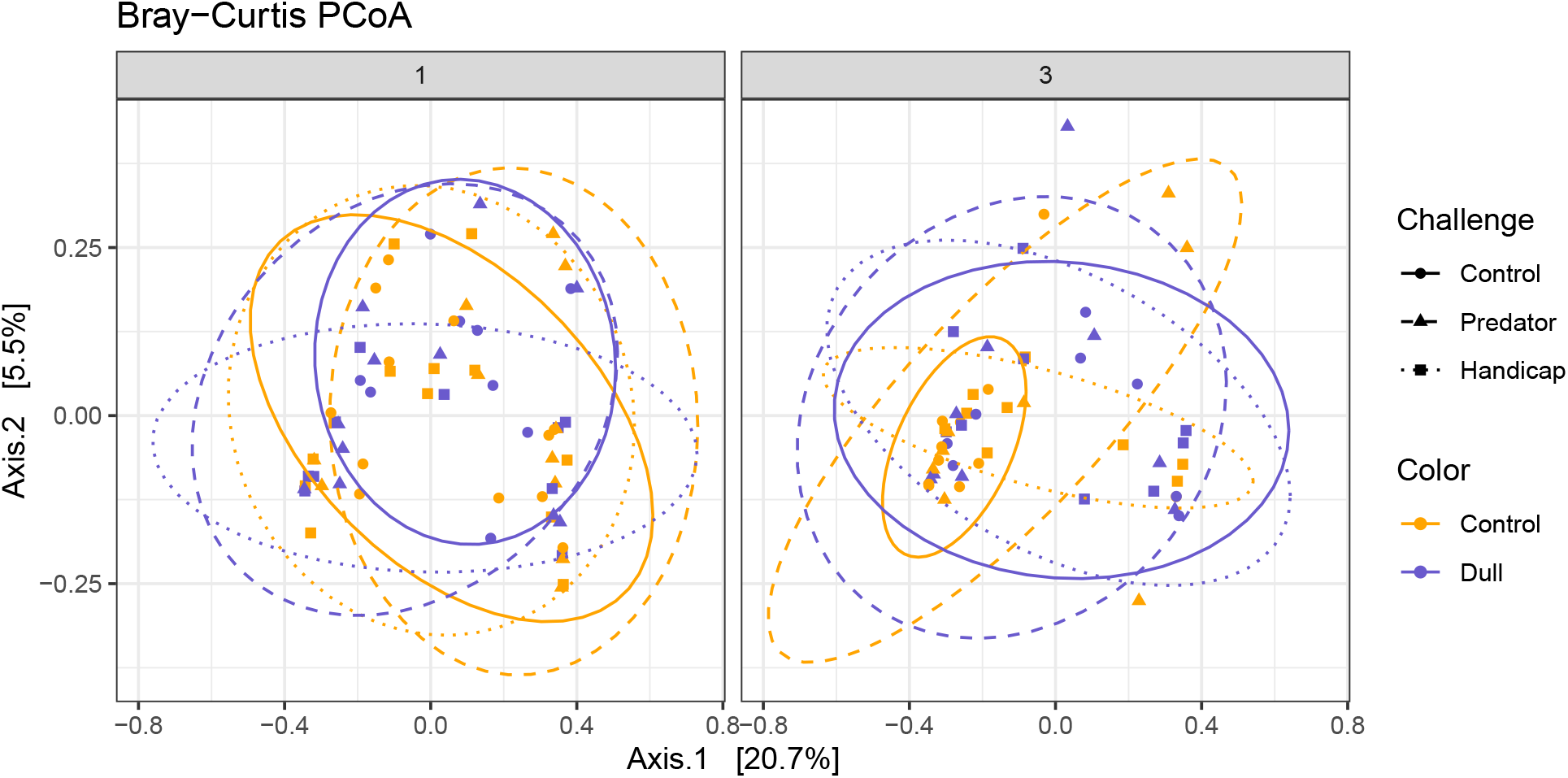
Bray-Curtis PCoA ordination plots by treatment group and capture number for experiment one.

**Figure S4:**
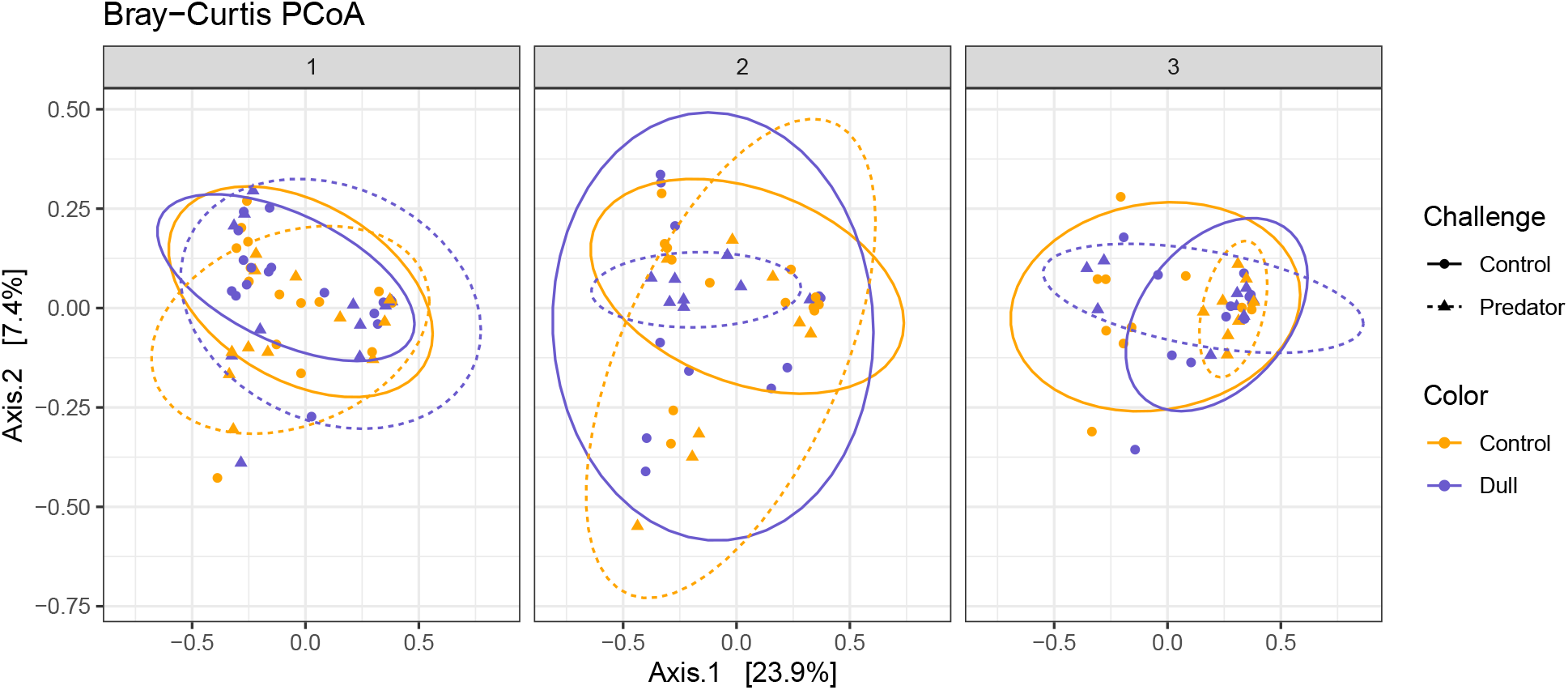
Bray-Curtis PCoA ordination plots by treatment group and capture number for experiment two.

**Table S1:** General linear model results for visits by other females and trips to other boxes by the focal female for each stage of experiment one. Models include nest box and day of year as random effects. Stage one models do not include challenge treatments because they had not occurred yet.

| Predictor | Estimate | CI | P |
| --- | --- | --- | --- |
| **Stage One: Visits by Other Females** |  |  |  |
| Intercept | -7.54 | -12.14 - -4.67 | < 0.001 |
| Signal Dulled | -0.57 | -4.60 - 3.10 | 0.72 |
| Day of Breeding | 0.10 | 0.01 - 0.21 | 0.02 |
| **Stage One: Trips by Focal Female** |  |  |  |
| Intercept | -7.08 | -11.82 - -2.36 | 0.003 |
| Signal Dulled | 1.22 | -1.01 - 3.46 | 0.28 |
| Day of Breeding | 0.58 | -0.15 - 1.30 | 0.12 |
| **Stage Two: Visits by Other Females** |  |  |  |
| Intercept | -0.70 | -1.71 - 0.31 | 0.17 |
| Signal Dulled | -0.17 | -1.54 - 1.20 | 0.81 |
| Predator | -1.04 | -2.46 - 0.37 | 0.15 |
| Handicap | -0.91 | -2.28 - 0.46 | 0.19 |
| Initial Brightness | 1.32 | 0.29 - 2.35 | 0.01 |
| Day of Breeding | 0.69 | 0.51 - 0.86 | < 0.001 |
| Dulled * Predator | 0.18 | -1.83 - 2.20 | 0.86 |
| Dulled * Handicap | -0.11 | -2.11 - 1.89 | 0.92 |
| Dulled * Brightness | -1.81 | -3.33 - -0.29 | 0.02 |
| Predator * Brightness | -1.73 | -3.04 - -0.42 | 0.01 |
| Handicap * Brightness | -1.99 | -3.69 - -0.30 | 0.02 |
| Dulled * Predator * Brightness | 2.60 | 0.53 - 4.68 | 0.01 |
| Dulled * Handicap * Brightness | 1.78 | -0.53 - 4.08 | 0.13 |
| **Stage Two: Trips by Focal Female** |  |  |  |
| Intercept | -1.84 | -2.64 - -1.04 | < 0.001 |
| Signal Dulled | -0.68 | -1.83 - 0.46 | 0.24 |
| Predator | -0.17 | -1.23 - 0.90 | 0.76 |
| Handicap | -0.82 | -1.96 - 0.33 | 0.16 |
| Day of Breeding | 0.66 | 0.45 - 0.87 | < 0.001 |
| Dulled * Predator | 0.66 | -0.93 - 2.25 | 0.42 |
| Dulled * Handicap | 1.51 | -0.13 - 3.14 | 0.07 |

**Table S2:** General linear model results for visits by other females and trips to other boxes by the focal female for each stage of experiment two. Models include nest box and day of year as random effects. Stage one models do not include dulling treatment because it had not occurred yet.

| Predictor | Estimate | CI | P |
| --- | --- | --- | --- |
| **Stage One: Visits by Other Females** |  |  |  |
| Intercept | -8.33 | -12.33 - -4.33 | < 0.001 |
| Predator | -0.25 | -4.95 - 4.45 | 0.92 |
| Day of Breeding | -0.04 | -0.14 - 0.07 | 0.51 |
| **Stage One: Trips by Focal Female** |  |  |  |
| Intercept | -8.73 | -13.69 - -3.77 | < 0.001 |
| Predator | 1.16 | -3.63 - 5.96 | 0.63 |
| Day of Breeding | 0.01 | -0.07 - 0.08 | 0.87 |
| **Stage Two: Visits by Other Females** |  |  |  |
| Intercept | -1.21 | -2.42 - 0.01 | 0.05 |
| Predator | 0.93 | -0.96 - 2.83 | 0.33 |
| Signal Dulled | -0.20 | -1.93 - 1.53 | 0.82 |
| Day of Breeding | 0.06 | 0.00 - 0.12 | 0.04 |
| Predator * Dulled | -0.28 | -2.94 - 2.37 | 0.84 |
| **Stage Two: Trips by Focal Female** |  |  |  |
| Intercept | -1.26 | -2.99 - 0.48 | 0.15 |
| Predator | -2.84 | -5.82 - 0.15 | 0.06 |
| Signal Dulled | -3.17 | -5.90 - -0.43 | 0.02 |
| Initial Brightness | -0.91 | -3.22 - 1.40 | 0.44 |
| Day of Breeding | 0.43 | 0.23 - 0.64 | < 0.001 |
| Predator * Dulled | 4.40 | 0.21 - 8.60 | 0.04 |
| Predator * Brightness | 1.90 | -1.57 - 5.36 | 0.28 |
| Dulled * Brightness | 2.00 | -0.93 - 4.93 | 0.18 |
| Predator * Dulled * Brightness | -4.46 | -8.79 - -0.13 | 0.04 |

**Table S3:** General linear model results for post-treatment mass in experiment one and experiment two. Both models also include female identity as a random effect.

| Predictor | Estimate | CI | P |
| --- | --- | --- | --- |
| **Experiment One: n = 57 birds; 110 measures** |  |  |  |
| Intercept (Control, 2nd Capture) | 20.03 | 19.28 - 20.77 | < 0.001 |
| Signal Dulled | 0.20 | -0.92 - 1.32 | 0.73 |
| Predator | -0.47 | -1.51 - 0.56 | 0.37 |
| Handicap | 0.49 | -0.59 - 1.58 | 0.37 |
| Stage (3rd Capture) | -1.53 | -2.49 - -0.57 | 0.002 |
| Pre-Treatment Mass | 0.60 | 0.36 - 0.85 | < 0.001 |
| Dulled * Predator | 0.38 | -1.14 - 1.91 | 0.63 |
| Dulled * Handicap | 0.07 | -1.51 - 1.65 | 0.93 |
| Dulled * Stage | -0.01 | -1.45 - 1.44 | 0.99 |
| Predator * Stage | 0.39 | -1.01 - 1.79 | 0.59 |
| Handicap * Stage | -0.14 | -1.54 - 1.26 | 0.85 |
| Dulled * Predator * Stage | -0.24 | -2.27 - 1.79 | 0.82 |
| Dulled * Handicap * Stage | 0.04 | -2.00 - 2.08 | 0.97 |
| **Experiment Two: n = 54 birds; 99 measures** |  |  |  |
| Intercept (Control, 2nd Capture) | 20.36 | 19.80 - 20.93 | < 0.001 |
| Signal Dulled | -0.34 | -1.14 - 0.46 | 0.41 |
| Predator | -0.79 | -1.64 - 0.06 | 0.07 |
| Stage (3rd Capture) | -2.02 | -2.72 - -1.32 | < 0.001 |
| Pre-Treatment Mass | 0.70 | 0.46 - 0.95 | < 0.001 |
| Dulled * Predator | 0.61 | -0.60 - 1.82 | 0.32 |
| Dulled * Stage | -0.23 | -1.22 - 0.77 | 0.66 |
| Predator * Stage | 0.71 | -0.37 - 1.80 | 0.20 |
| Dulled * Predator * Stage | -0.28 | -1.80 - 1.24 | 0.72 |

**Table S4:** Results of linear models for each corticosterone measurement recorded in experiment one and experiment two. Second capture models only include the first stage of treatment because the second stage had not occurred yet.

| Predictor | Estimate | CI | P |
| --- | --- | --- | --- |
| **Experiment One: 2nd Capture Base Cort** |  |  |  |
| Intercept | 2.40 | 1.50 - 3.30 | < 0.001 |
| Signal Dulled | -0.11 | -1.41 - 1.18 | 0.86 |
| Initial Base Cort | -0.15 | -0.77 - 0.47 | 0.62 |
| **Experiment One: 3rd Capture Base Cort** |  |  |  |
| Intercept | 1.93 | -0.49 - 4.34 | 0.12 |
| Signal Dulled | 1.64 | -2.00 - 5.29 | 0.37 |
| Initial Base Cort | 0.11 | -0.89 - 1.12 | 0.82 |
| Predator | 0.50 | -3.12 - 4.12 | 0.78 |
| Handicap | 1.62 | -1.88 - 5.12 | 0.36 |
| Dulled * Predator | 0.06 | -5.13 - 5.26 | 0.98 |
| Duleld * Handicap | -1.15 | -6.33 - 4.03 | 0.66 |
| **Experiment One: 3rd Capture Induced Cort** |  |  |  |
| Intercept | 30.64 | 21.19 - 40.08 | < 0.001 |
| Signal Dulled | -3.56 | -17.63 - 10.51 | 0.61 |
| Initial Induced Cort | 7.58 | 3.64 - 11.52 | < 0.001 |
| Predator | 2.07 | -12.03 - 16.17 | 0.77 |
| Handicap | -0.82 | -14.52 - 12.88 | 0.91 |
| Dulled * Predator | 8.28 | -11.84 - 28.40 | 0.41 |
| Dulled * Handicap | 5.23 | -14.61 - 25.06 | 0.60 |
| **Experiment One: 3rd Capture Post-Dex Cort** |  |  |  |
| Intercept | 13.67 | 10.30 - 17.04 | < 0.001 |
| Signal Dulled | 0.74 | -4.27 - 5.74 | 0.77 |
| Initial Post-Dex Cort | 3.42 | 1.85 - 4.98 | < 0.001 |
| Predator | 3.28 | -1.74 - 8.30 | 0.20 |
| Handicap | -1.53 | -6.49 - 3.44 | 0.54 |
| Dulled * Predator | -4.47 | -11.85 - 2.91 | 0.23 |
| Dulled * Handicap | 0.44 | -6.65 - 7.53 | 0.90 |
| **Experiment Two: 2nd Capture Base Cort** |  |  |  |
| Intercept | 2.46 | 1.49 - 3.42 | < 0.001 |
| Predator | 0.63 | -0.82 - 2.09 | 0.39 |
| Initial Base Cort | 1.13 | 0.40 - 1.85 | 0.003 |
| **Experiment Two: 3rd Capture Base Cort** |  |  |  |
| Intercept | 2.22 | 0.85 - 3.60 | 0.002 |
| Predator | 0.55 | -1.56 - 2.65 | 0.60 |
| Initial Base Cort | 0.03 | -0.69 - 0.76 | 0.92 |
| Signal Dulled | 0.78 | -1.13 - 2.70 | 0.41 |
| Dulled * Predator | -1.66 | -4.58 - 1.25 | 0.26 |

**Table S4:** Results of linear models for each corticosterone measurement recorded in experiment one and experiment two. Second capture models only include the first stage of treatment because the second stage had not occurred yet. (*continued*)
| Predictor | Estimate | CI | P |
| --- | --- | --- | --- |
| **Experiment Two: 3rd Capture Induced Cort** |  |  |  |
| Intercept | 28.72 | 20.92 - 36.52 | < 0.001 |
| Initial Induced Cort | 4.50 | 0.17 - 8.84 | 0.04 |
| Signal Dulled | 1.48 | -10.20 - 13.15 | 0.80 |
| Dulled * Predator | -0.43 | -17.91 - 17.04 | 0.96 |
| **Experiment Two: 3rd Capture Post-ACTH Cort** |  |  |  |
| Intercept | 39.32 | 25.73 - 52.90 | < 0.001 |
| Predator | -0.73 | -21.97 - 20.52 | 0.95 |
| Signal Dulled | 2.49 | -16.72 - 21.71 | 0.80 |
| Dulled * Predator | 6.87 | -22.73 - 36.47 | 0.64 |

**Table S5:**
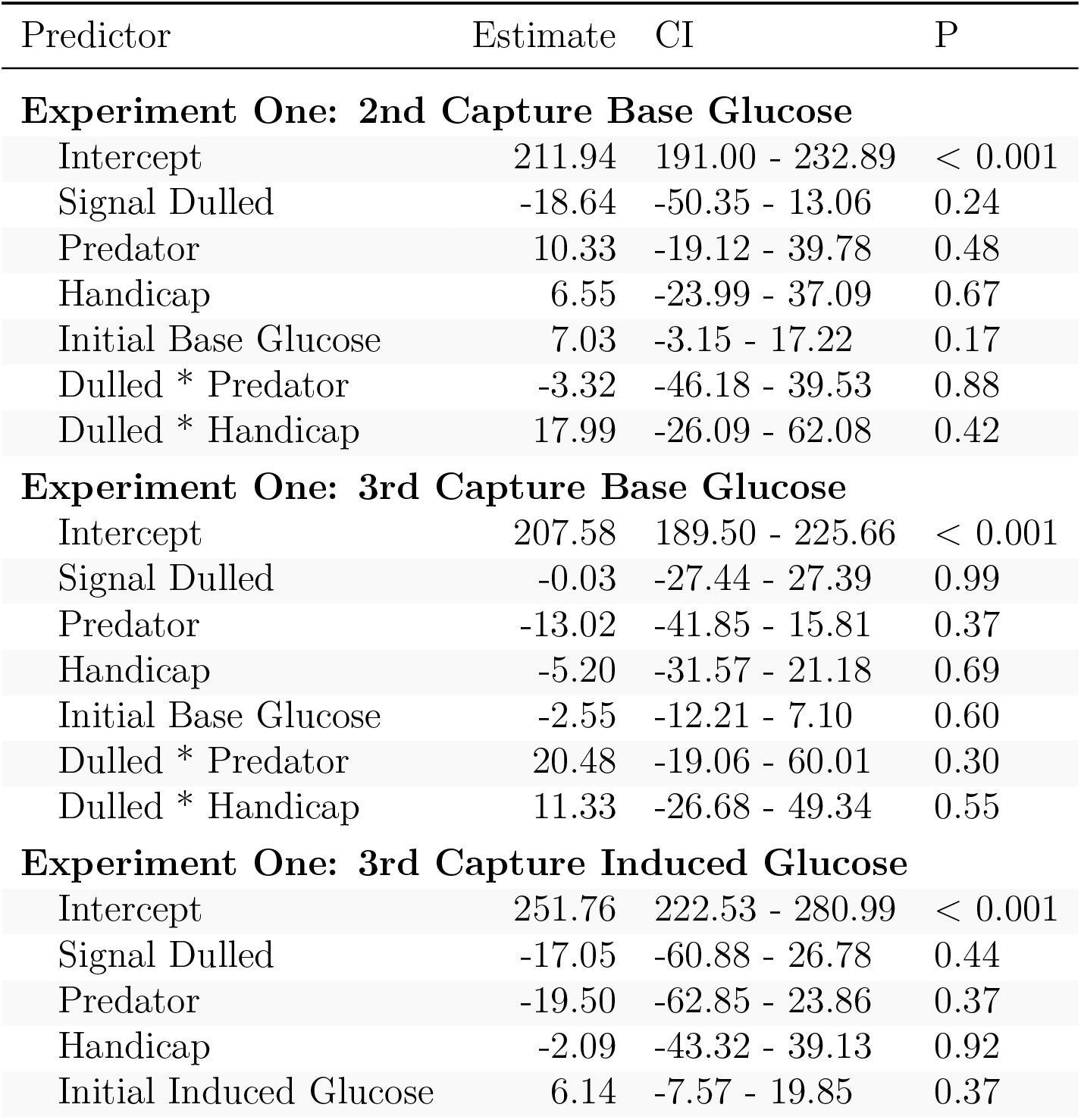
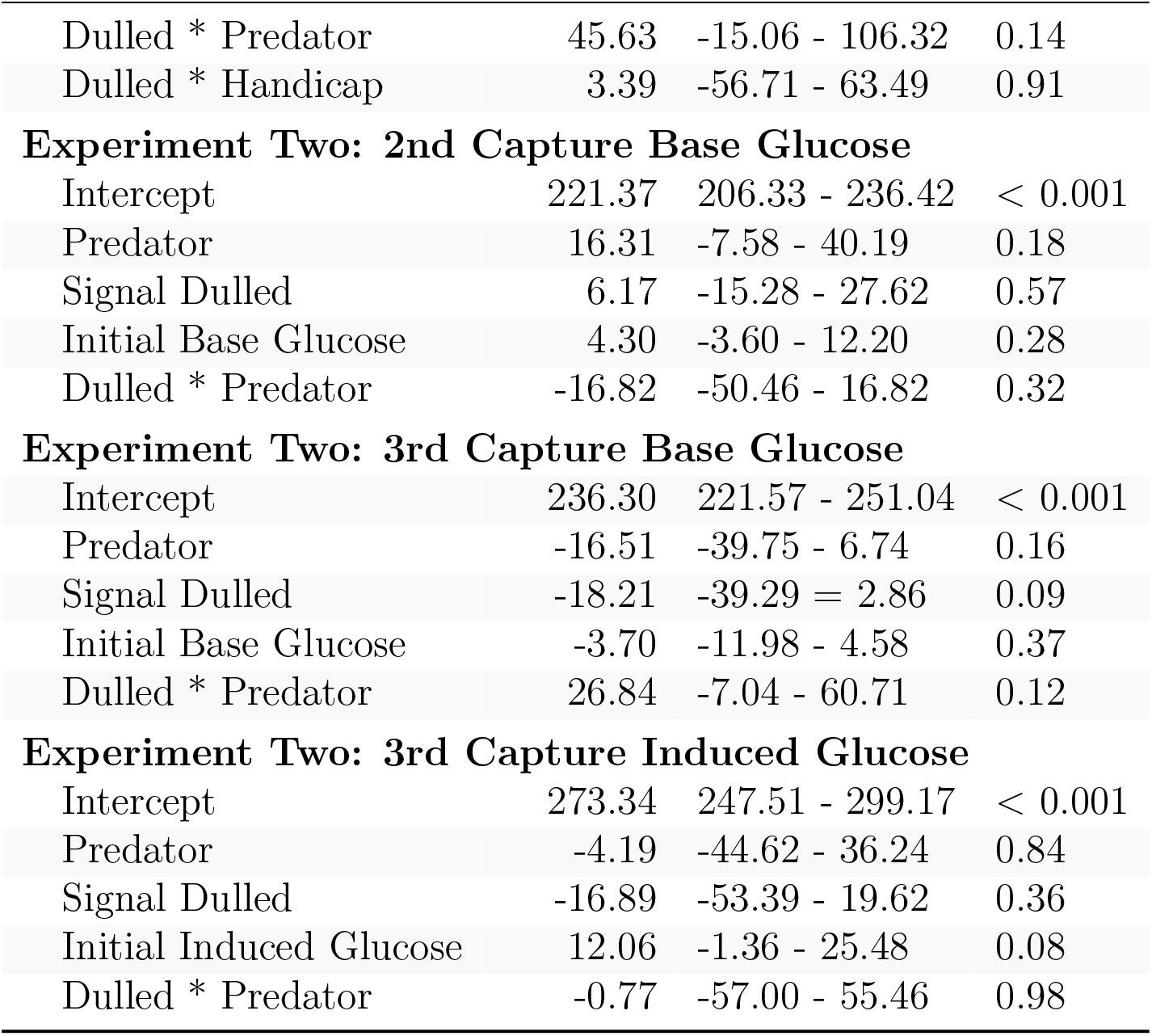
Results of linear models for each glucose measurement recorded in experiment one and experiment two. Second capture models only include the first stage of treatment because the second stage had not occurred yet.

| Predictor | Estimate | CI | P |
| --- | --- | --- | --- |
| **Experiment One: 2nd Capture Base Glucose** |  |  |  |
| Intercept | 211.94 | 191.00 - 232.89 | < 0.001 |
| Signal Dulled | -18.64 | -50.35 - 13.06 | 0.24 |
| Predator | 10.33 | -19.12 - 39.78 | 0.48 |
| Handicap | 6.55 | -23.99 - 37.09 | 0.67 |
| Initial Base Glucose | 7.03 | -3.15 - 17.22 | 0.17 |
| Dulled * Predator | -3.32 | -46.18 - 39.53 | 0.88 |
| Dulled * Handicap | 17.99 | -26.09 - 62.08 | 0.42 |
| **Experiment One: 3rd Capture Base Glucose** |  |  |  |
| Intercept | 207.58 | 189.50 - 225.66 | < 0.001 |
| Signal Dulled | -0.03 | -27.44 - 27.39 | 0.99 |
| Predator | -13.02 | -41.85 - 15.81 | 0.37 |
| Handicap | -5.20 | -31.57 - 21.18 | 0.69 |
| Initial Base Glucose | -2.55 | -12.21 - 7.10 | 0.60 |
| Dulled * Predator | 20.48 | -19.06 - 60.01 | 0.30 |
| Dulled * Handicap | 11.33 | -26.68 - 49.34 | 0.55 |
| **Experiment One: 3rd Capture Induced Glucose** |  |  |  |
| Intercept | 251.76 | 222.53 - 280.99 | < 0.001 |
| Signal Dulled | -17.05 | -60.88 - 26.78 | 0.44 |
| Predator | -19.50 | -62.85 - 23.86 | 0.37 |
| Handicap | -2.09 | -43.32 - 39.13 | 0.92 |
| Initial Induced Glucose | 6.14 | -7.57 - 19.85 | 0.37 |

**Table S5:** Results of linear models for each glucose measurement recorded in experiment one and experiment two. Second capture models only include the first stage of treatment because the second stage had not occurred yet. (*continued*)
| Predictor | Estimate | CI | P |
| --- | --- | --- | --- |
| Dulled * Predator | 45.63 | -15.06 - 106.32 | 0.14 |
| Dulled * Handicap | 3.39 | -56.71 - 63.49 | 0.91 |
| **Experiment Two: 2nd Capture Base Glucose** |  |  |  |
| Intercept | 221.37 | 206.33 - 236.42 | < 0.001 |
| Predator | 16.31 | -7.58 - 40.19 | 0.18 |
| Signal Dulled | 6.17 | -15.28 - 27.62 | 0.57 |
| Initial Base Glucose | 4.30 | -3.60 - 12.20 | 0.28 |
| Dulled * Predator | -16.82 | -50.46 - 16.82 | 0.32 |
| **Experiment Two: 3rd Capture Base Glucose** |  |  |  |
| Intercept | 236.30 | 221.57 - 251.04 | < 0.001 |
| Predator | -16.51 | -39.75 - 6.74 | 0.16 |
| Signal Dulled | -18.21 | -39.29 - 2.86 | 0.09 |
| Initial Base Glucose | -3.70 | -11.98 - 4.58 | 0.37 |
| Dulled * Predator | 26.84 | -7.04 - 60.71 | 0.12 |
| **Experiment Two: 3rd Capture Induced Glucose** |  |  |  |
| Intercept | 273.34 | 247.51 - 299.17 | < 0.001 |
| Predator | -4.19 | -44.62 - 36.24 | 0.84 |
| Signal Dulled | -16.89 | -53.39 - 19.62 | 0.36 |
| Initial Induced Glucose | 12.06 | -1.36 - 25.48 | 0.08 |
| Dulled * Predator | -0.77 | -57.00 - 55.46 | 0.98 |

**Table S6:** Results of linear models for alpha diversity at each capture measured as Faith’s phylogenetic diversity (PD), Shannon Index, or Inverse Simpson in each experiment. Second capture models only include the first stage of treatment because the second stage had not occurred yet.

| Predictor | Estimate | CI | P |
| --- | --- | --- | --- |
| **Experiment One: 3rd Capture PD** |  |  |  |
| Intercept | 5.75 | 3.25 - 8.24 | < 0.001 |
| Pre-treatment Diversity | 0.02 | -0.20 - 0.24 | 0.86 |
| Signal Dulled | 1.45 | -1.41 - 4.32 | 0.31 |
| Predator | 0.50 | -2.34 - 3.35 | 0.72 |
| Handicap | 1.26 | -1.51 - 4.04 | 0.37 |
| Dulled * Predator | -1.95 | -6.02 - 2.12 | 0.34 |
| Dulled * Handicap | -1.06 | -5.08 - 2.96 | 0.60 |
| **Experiment One: 3rd Capture Shannon** |  |  |  |
| Intercept | 1.00 | 0.33 - 1.67 | 0.004 |
| Pre-treatment Diversity | 0.11 | -0.15 - 0.36 | 0.41 |
| Signal Dulled | 0.59 | -0.18 - 1.37 | 0.13 |

**Table S6:** Results of linear models for alpha diversity at each capture measured as Faith's phylogenetic diversity (PD), Shannon Index, or Inverse Simpson in each experiment.
| Predictor | Estimate | CI | P |
| --- | --- | --- | --- |
| Predator | 0.42 | -0.36 - 1.19 | 0.28 |
| Handicap | 0.82 | 0.07 - 1.57 | 0.03 |
| Dulled * Predator | -0.85 | -1.97 - 0.27 | 0.13 |
| Dulled * Handicap | -0.64 | -1.73 - 0.45 | 0.24 |
| **Experiment One: 3rd Capture Inverse Simpson** |  |  |  |
| Intercept | 2.06 | -0.81 - 4.93 | 0.16 |
| Pre-treatment Diversity | -0.06 | -0.22 - 0.35 | 0.66 |
| Signal Dulled | 4.89 | 0.85 - 8.94 | 0.02 |
| Predator | 1.13 | -2.93 - 5.20 | 0.58 |
| Handicap | 2.34 | -1.56 - 6.24 | 0.23 |
| Dulled * Predator | -5.33 | -11.16 - 0.51 | 0.07 |
| Dulled * Handicap | -4.00 | -9.69 - 1.68 | 0.16 |
| **Experiment Two: 2nd Capture PD** |  |  |  |
| Intercept | 5.26 | 2.44 - 8.08 | 0.001 |
| Pre-treatment Diversity | 0.04 | -0.28 - 0.36 | 0.80 |
| Signal Dulled | 0.96 | -1.24 - 3.16 | 0.38 |
| Predator | 0.32 | -2.34 - 2.98 | 0.81 |
| Dulled * Predator | -0.52 | -4.33 - 3.28 | 0.78 |
| **Experiment Two: 2nd Capture Shannon** |  |  |  |
| Intercept | 0.90 | 0.32 - 1.48 | 0.004 |
| Pre-treatment Diversity | 0.31 | 0.03 - 0.59 | 0.03 |
| Signal Dulled | 0.06 | -0.45 - 0.57 | 0.81 |
| Predator | -0.06 | -0.69 - 0.57 | 0.84 |
| Dulled * Predator | 0.64 | -0.25 - 1.53 | 0.15 |
| **Experiment Two: 2nd Capture Inverse Simpson** |  |  |  |
| Intercept | 2.42 | 0.50 - 4.34 | 0.02 |
| Pre-treatment Diversity | 0.20 | -0.06 - 0.45 | 0.13 |
| Signal Dulled | 0.47 | -1.83 - 2.76 | 0.21 |
| Predator | -0.28 | -3.14 - 2.58 | 0.84 |
| Dulled * Predator | 2.11 | -1.91 - 6.14 | 0.29 |
| **Experiment Two: 3rd Capture PD** |  |  |  |
| Intercept | 4.95 | 0.35 - 9.55 | 0.04 |
| Pre-treatment Diversity | 0.45 | -0.05 - 0.94 | 0.08 |
| Signal Dulled | 0.21 | -3.01 - 3.42 | 0.90 |
| Predator | 1.45 | -2.42 - 5.33 | 0.45 |
| Dulled * Predator | -2.60 | -7.77 - 2.56 | 0.31 |
| **Experiment Two: 3rd Capture Shannon** |  |  |  |
| Intercept | 1.33 | 0.46 - 2.20 | 0.004 |

**Table S6:** Results of linear models for alpha diversity at each capture measured as Faith's phylogenetic diversity (PD), Shannon Index, or Inverse Simpson in each experiment.
| Predictor | Estimate | CI | P |
| --- | --- | --- | --- |
| Pre-treatment Diversity | 0.24 | -0.14 - 0.61 | 0.20 |
| Signal Dulled | 0.00 | -0.73 - 0.73 | 0.99 |
| Predator | -0.39 | -1.27 - 0.48 | 0.36 |
| Dulled * Predator | 0.20 | -0.98 - 1.38 | 0.74 |
| **Experiment Two: 3rd Capture Inverse Simpson** |  |  |  |
| Intercept | 2.93 | 0.48 - 5.38 | 0.02 |
| Pre-treatment Diversity | 0.31 | 0.03 - 0.60 | 0.03 |
| Signal Dulled | -2.59 | -5.96 - 0.77 | 0.13 |
| Predator | -0.27 | -3.06 - 2.53 | 0.85 |
| Dulled * Predator | 2.42 | -2.10 - 6.94 | 0.28 |

**Table S7:**
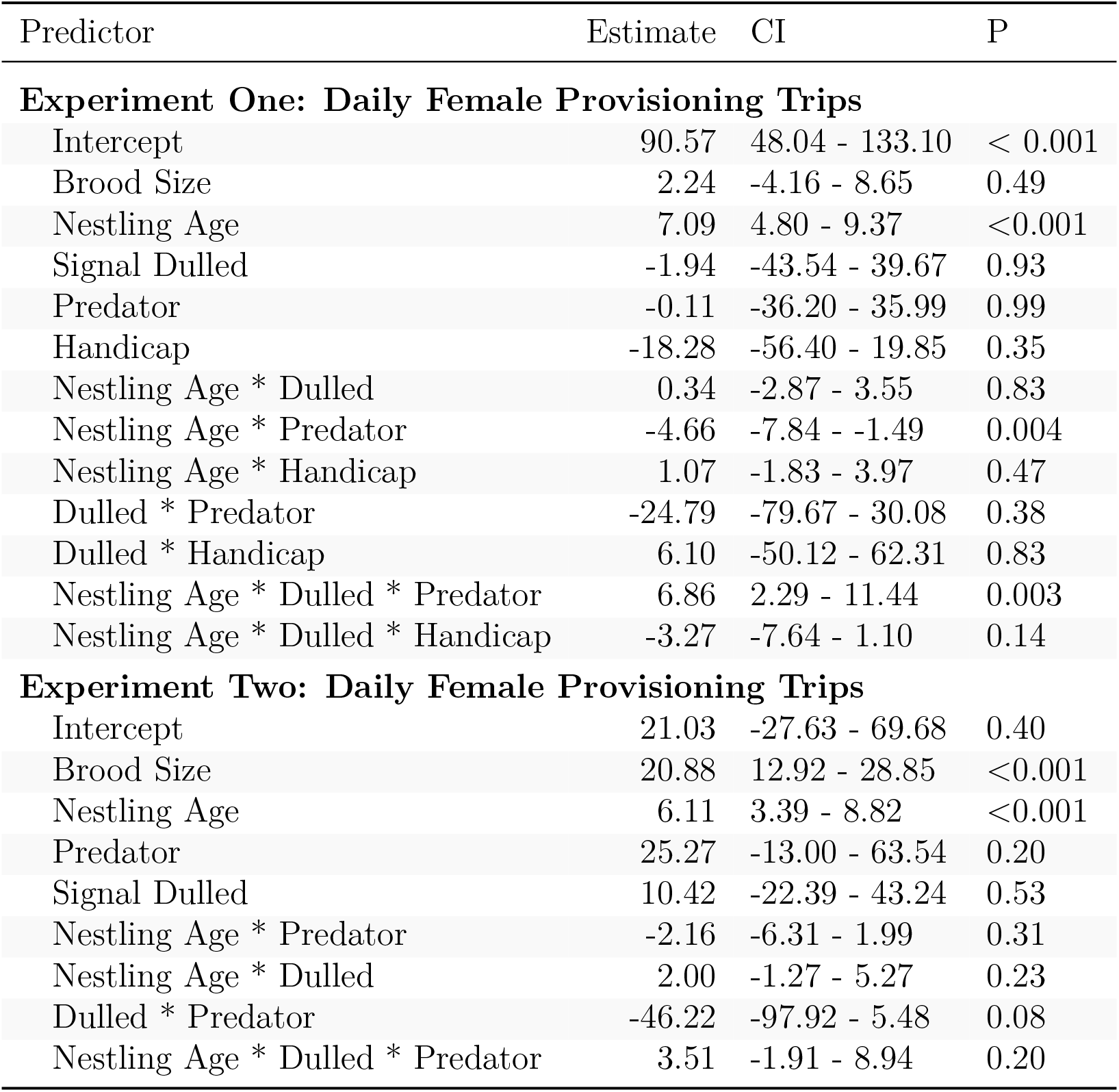
Results of GLMMs with total daily provisioning by females as the response variable for each experiment. Models include female identity and day of year as random effects with a Poisson distribution.

**Table S8:**
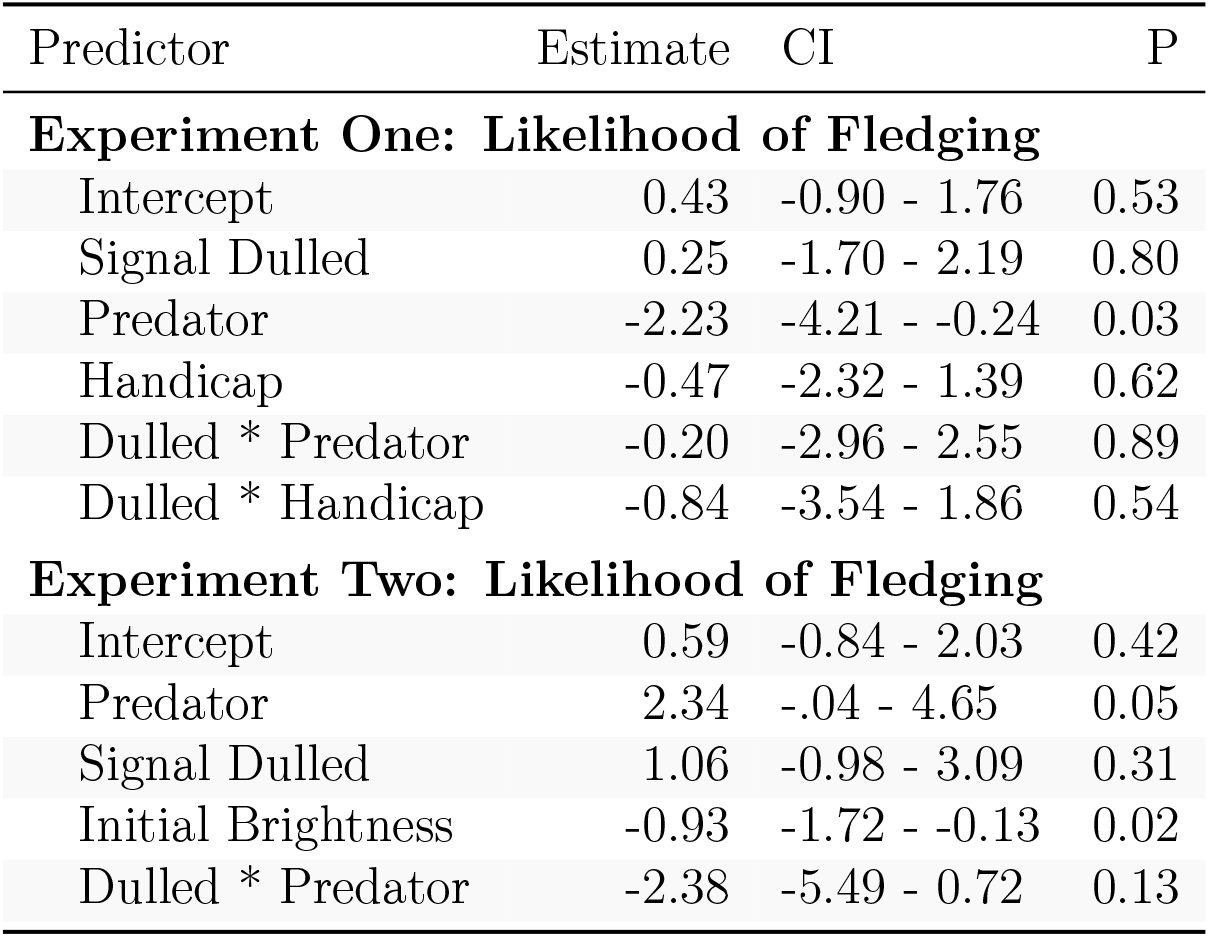
Results of binomial GLMMs for fledging success in each year. Models include nest identity as a random effect.

| Predictor | Estimate | CI | P |
| --- | --- | --- | --- |
| **Experiment One: Likelihood of Fledging** |  |  |  |
| Intercept | 0.43 | -0.90 - 1.76 | 0.53 |
| Signal Dulled | 0.25 | -1.70 - 2.19 | 0.80 |
| Predator | -2.23 | -4.21 - -0.24 | 0.03 |
| Handicap | -0.47 | -2.32 - 1.39 | 0.62 |
| Dulled * Predator | -0.20 | -2.96 - 2.55 | 0.89 |
| Dulled * Handicap | -0.84 | -3.54 - 1.86 | 0.54 |
| **Experiment Two: Likelihood of Fledging** |  |  |  |
| Intercept | 0.59 | -0.84 - 2.03 | 0.42 |
| Predator | 2.34 | -.04 - 4.65 | 0.05 |
| Signal Dulled | 1.06 | -0.98 - 3.09 | 0.31 |
| Initial Brightness | -0.93 | -1.72 - -0.13 | 0.02 |
| Dulled * Predator | -2.38 | -5.49 - 0.72 | 0.13 |

## REFERENCES

Anderson, J. A., Johnston, R. A., Lea, A. J., Campos, F. A., Voyles, T. N., Akinyi, M. Y., Alberts, S. C., Archie, E. A., & Tung, J. (2021). High social status males experience accelerated epigenetic aging in wild baboons. Elife, 10, e66128.

Anderson, J. A., Lea, A. J., Voyles, T. N., Akinyi, M. Y., Nyakundi, R., Ochola, L., Omondi, M., Nyundo, F., Zhang, Y., Campos, F. A. others. (2022). Distinct gene regulatory signatures of dominance rank and social bond strength in wild baboons. Philosophical Transactions of the Royal Society B, 377 (1845), 20200441.

Bailey, M. T., Dowd, S. E., Galley, J. D., Hufnagle, A. R., Allen, R. G., & Lyte, M. (2011). Exposure to a social stressor alters the structure of the intestinal microbiota: Implications for stressor-induced immunomodulation. Brain, Behavior, and Immunity, 25 (3), 397–407.

Bates, D., Mächler, M., Bolker, B., & Walker, S. (2014). Fitting linear mixed-effects models using lme4. arXiv Preprint arXiv:1406.5823.

Beck, M. L., Hopkins, W. A., & Hawley, D. M. (2015). Relationships among plumage coloration, blood selenium concentrations and immune responses of adult and nestling tree swallows. Journal of Experimental Biology, 218 (21), 3415–3424.

Berzins, L. L., & Dawson, R. D. (2016). Experimentally altered plumage brightness of female tree swallows: A test of the differential allocation hypothesis. Behaviour, 153 (5), 525–550.

Berzins, L. L., & Dawson, R. D. (2018). Experimentally altered plumage brightness of female tree swallows (Tachycineta bicolor) influences nest site retention and reproductive success. Canadian Journal of Zoology, 96 (6), 600–607.

Callahan, B. J., McMurdie, P. J., Rosen, M. J., Han, A. W., Johnson, A. J. A., & Holmes, S. P. (2016). DADA2: High-resolution sample inference from Illumina amplicon data. Nature Methods, 13 (7), 581–583.

Callahan, B. J., Sankaran, K., Fukuyama, J. A., McMurdie, P. J., & Holmes, S. P. (2016). Bioconductor workflow for microbiome data analysis: From raw reads to community analyses. F1000Research, 5.

Caporaso, J. G., Lauber, C. L., Walters, W. A., Berg-Lyons, D., Huntley, J., Fierer, N., Owens, S. M., Betley, J., Fraser, L., & Bauer, M. (2012). Ultra-high-throughput microbial community analysis on the Illumina HiSeq and MiSeq platforms. The ISME Journal, 6 (8), 1621–1624.

Caporaso, J. G., Lauber, C. L., Walters, W. A., Berg-Lyons, D., Lozupone, C. A., Turnbaugh, P. J., Fierer, N., & Knight, R. (2011). Global patterns of 16S rRNA diversity at a depth of millions of sequences per sample. Proceedings of the National Academy of Sciences, 108 (Supplement 1), 4516–4522.

Charuvastra, A., & Cloitre, M. (2008). Social bonds and posttraumatic stress disorder. Annual Review of Psychology, 59, 301.

Clinchy, M., Sheriff, M. J., & Zanette, L. Y. (2013). Predator-induced stress and the ecology of fear. Functional Ecology, 27 (1), 56–65.

Coady, C. D., & Dawson, R. D. (2013). Subadult plumage color of female tree swallows (Tachycineta bicolor) reduces conspecific aggression during the breeding season. The Wilson Journal of Ornithology, 125 (2), 348–357.

Davis, N. M., Proctor, D. M., Holmes, S. P., Relman, D. A., & Callahan, B. J. (2018). Simple statistical identification and removal of contaminant sequences in marker-gene and metagenomics data. Microbiome, 6 (1), 226.

Dixon, P. (2003). VEGAN, a package of r functions for community ecology. Journal of Vegetation Science, 14 (6), 927–930.

Holt-Lunstad, J., Smith, T. B., & Layton, J. B. (2010). Social relationships and mortality risk: A meta-analytic review. PLoS Medicine, 7 (7), e1000316.

Houtz, J. L., Taff, C. C., & Vitousek, M. N. (2022). Gut microbiome as a mediator of stress resilience: A reactive scope model framework. Integrative and Comparative Biology.

Ibáñez-Álamo, J., Chastel, O., & Soler, M. (2011). Hormonal response of nestlings to predator calls. General and Comparative Endocrinology, 171 (2), 232–236.

Kembel, S. W., Cowan, P. D., Helmus, M. R., Cornwell, W. K., Morlon, H., Ackerly, D. D., Blomberg, S. P., & Webb, C. O. (2010). Picante: R tools for integrating phylogenies and ecology. Bioinformatics, 26 (11), 1463–1464.

Kuznetsova, A., Brockhoff, P. B., & Christensen, R. H. (2017). lmerTest package: Tests in linear mixed effects models. Journal of Statistical Software, 82, 1–26.

LaManna, J. A., & Martin, T. E. (2016). Costs of fear: Behavioural and life-history responses to risk and their demographic consequences vary across species. Ecology Letters, 19 (4), 403–413.

Lenth, R. V. (2020). Emmeans: Estimated marginal means, aka least-squares means. https://CRAN.R-project.org/package=emmeans

Levin, I. I., Zonana, D. M., Fosdick, B. K., Song, S. J., Knight, R., & Safran, R. J. (2016). Stress response, gut microbial diversity and sexual signals correlate with social interactions. Biology Letters, 12 (6), 20160352.

Lima, S. L. (2009). Predators and the breeding bird: Behavioral and reproductive flexibility under the risk of predation. Biological Reviews, 84 (3), 485–513.

Lupien, S. J., McEwen, B. S., Gunnar, M. R., & Heim, C. (2009). Effects of stress throughout the lifespan on the brain, behaviour and cognition. Nature Reviews Neuroscience, 10 (6), 434–445.

Maia, R., Eliason, C. M., Bitton, P., Doucet, S. M., & Shawkey, M. D. (2013). Pavo: An R package for the analysis, visualization and organization of spectral data. Methods in Ecology and Evolution, 4 (10), 906–913.

McMurdie, P. J., & Holmes, S. (2013). Phyloseq: An r package for reproducible interactive analysis and graphics of microbiome census data. PloS One, 8 (4), e61217.

McNew, S. M., Taff, C. C., Zimmer, C., Uehling, J. J., Ryan, T. A., Chang van Oordt, D., Houtz, J. L., Injaian, A. S., & Vitousek, M. N. (2022). Developmental stage-dependent effects of perceived predation risk on physiology and fledging success of tree swallows (Tachycineta bicolor). BioRxiv.

Oksanen, J., Blanchet, F. G., Kindt, R., Legendre, P., Minchin, P., O’hara, R., Simpson, G., Solymos, P., Stevens, M. H. H., Wagner, H. others. (2013). Community ecology package. R Package Version, 2 (0), 321–326.

Quast, C., Pruesse, E., Yilmaz, P., Gerken, J., Schweer, T., Yarza, P., Peplies, J., & Glöckner, F. O. (2012). The SILVA ribosomal RNA gene database project: Improved data processing and web-based tools. Nucleic Acids Research, 41 (D1), D590–D596.

R Core Team. (2020). R: A language and environment for statistical computing. R Foundation for Statistical Computing. https://www.R-project.org/

Rea, K., Dinan, T. G., & Cryan, J. F. (2016). The microbiome: A key regulator of stress and neuroinflammation. Neurobiology of Stress, 4, 23–33.

Schliep, K. P. (2011). Phangorn: Phylogenetic analysis in r. Bioinformatics, 27 (4), 592–593.

Senar, J. C., Domènech, J., & Uribe, F. (2002). Great tits (Parus major) reduce body mass in response to wing area reduction: A field experiment. Behavioral Ecology, 13 (6), 725–727.

Shipley, J. R., Twining, C. W., Taff, C. C., Vitousek, M. N., Flack, A., & Winkler, D. W. (2020). Birds advancing lay dates with warming springs face greater risk of chick mortality. Proceedings of the National Academy of Sciences, 117 (41), 25590–25594.

Shipley, J. R., Twining, C. W., Taff, C. C., Vitousek, M. N., & Winkler, D. W. (2022). Selection counteracts developmental plasticity in body-size responses to climate change. Nature Climate Change.

Snyder-Mackler, N., Burger, J. R., Gaydosh, L., Belsky, D. W., Noppert, G. A., Campos, F. A., Bartolomucci, A., Yang, Y. C., Aiello, A. E., O’Rand, A. others. (2020). Social determinants of health and survival in humans and other animals. Science, 368 (6493), eaax9553.

Snyder-Mackler, N., Sanz, J., Kohn, J. N., Brinkworth, J. F., Morrow, S., Shaver, A. O., Grenier, J.-C., Pique-Regi, R., Johnson, Z. P., Wilson, M. E. others. (2016). Social status alters immune regulation and response to infection in macaques. Science, 354 (6315), 1041–1045.

Snyder-Mackler, N., Sanz, J., Kohn, J. N., Voyles, T., Pique-Regi, R., Wilson, M. E., Barreiro, L. B., & Tung, J. (2019). Social status alters chromatin accessibility and the gene regulatory response to glucocorticoid stimulation in rhesus macaques. Proceedings of the National Academy of Sciences, 116 (4), 1219–1228.

Taff, C. C., Campagna, L., & Vitousek, M. N. (2019). Genome-wide variation in DNA methylation is associated with stress resilience and plumage brightness in a wild bird. Molecular Ecology, 28 (16), 3722–3737.

Taff, C. C., Johnson, B. A., Anker, A. T., Rodriguez, A. M., Houtz, J. L., Uehling, J. J., & Vitousek, M. N. (2022). No apparent trade-off between the quality of nest-grown feathers and time spent in the nest in an aerial insectivore, the tree swallow. Ornithology.

Taff, C. C., & Vitousek, M. N. (2016). Endocrine flexibility: Optimizing phenotypes in a dynamic world? Trends in Ecology & Evolution, 31 (6), 476–488.

Taff, C. C., Zimmer, C., Ryan, T. A., Oordt, D. C. van, Aborn, D. A., Ardia, D. R., Johnson, L. S., Rose, A. P., & Vitousek, M. N. (2022). Individual variation in natural or manipulated corticosterone does not covary with circulating glucose in a wild bird. Journal of Experimental Biology, 225 (4), jeb243262.

Taff, C. C., Zimmer, C., Scheck, D., Ryan, T. A., Houtz, J. L., Smee, M. R., Hendry, T. A., & Vitousek, M. N. (2021). Plumage manipulation alters associations between behaviour, physiology, the internal microbiome and fitness. Animal Behaviour, 178, 11–36.

Taff, C. C., Zimmer, C., & Vitousek, M. N. (2019). Achromatic plumage brightness predicts stress resilience and social interactions in tree swallows (Tachycineta bicolor). Behavioral Ecology, 30 (3), 733–745.

Tung, J., Barreiro, L. B., Burns, M. B., Grenier, J.-C., Lynch, J., Grieneisen, L. E., Altmann, J., Alberts, S. C., Blekhman, R., & Archie, E. A. (2015). Social networks predict gut microbiome composition in wild baboons. Elife, 4, e05224.

Uehling, J. J., Taff, C. C., Winkler, D. W., & Vitousek, M. N. (2020). Developmental temperature predicts the adult response to stressors in a free-living passerine. Journal of Animal Ecology, 89 (3), 842–854.

Vitousek, M. N., Taff, C. C., Ardia, D. R., Stedman, J. M., Zimmer, C., Salzman, T. C., & Winkler, D. W. (2018). The lingering impact of stress: Brief acute glucocorticoid exposure has sustained, dose-dependent effects on reproduction. Proceedings of the Royal Society B: Biological Sciences, 285 (1882), 20180722.

Vitousek, M. N., Taff, C. C., Hallinger, K. K., Zimmer, C., & Winkler, D. W. (2018). Hormones and fitness: Evidence for trade-offs in glucocorticoid regulation across contexts. Frontiers in Ecology and Evolution, 6, 42.

Vo, A., & Jedlicka, J. A. (2014). Protocols for metagenomic DNA extraction and Illumina amplicon library preparation for faecal and swab samples. Molecular Ecology Resources, 14 (6), 1183–1197.

Winkler, D. W., & Allen, P. E. (1995). Effects of handicapping on female condition and reproduction in tree swallows (Tachycineta bicolor). The Auk, 112 (3), 737–747.

Winkler, D. W., Hallinger, K. K., Pegan, T. M., Taff, C. C., Verhoeven, M. A., Chang van Oordt, D., Stager, M., Uehling, J. J., Vitousek, M. N., Andersen, M. J. others. (2020). Full lifetime perspectives on the costs and benefits of lay-date variation in tree swallows. Ecology, 101 (9), e03109.

Winkler, D. W., Luo, M. K., & Rakhimberdiev, E. (2013). Temperature effects on food supply and chick mortality in tree swallows (tachycineta bicolor). Oecologia, 173 (1), 129–138.

Yang, Y. C., Boen, C., Gerken, K., Li, T., Schorpp, K., & Harris, K. M. (2016). Social relationships and physiological determinants of longevity across the human life span. Proceedings of the National Academy of Sciences, 113 (3), 578–583.

Yilmaz, P., Parfrey, L. W., Yarza, P., Gerken, J., Pruesse, E., Quast, C., Schweer, T., Peplies, J., Ludwig, W., & Glöckner, F. O. (2014). The SILVA and “all-species living tree project (LTP)” taxonomic frameworks. Nucleic Acids Research, 42 (D1), D643–D648.

Zanette, L. Y., White, A. F., Allen, M. C., & Clinchy, M. (2011). Perceived predation risk reduces the number of offspring songbirds produce per year. Science, 334 (6061), 1398–1401.

Zimmer, C., Taff, C. C., Ardia, D. R., Ryan, T. A., Winkler, D. W., & Vitousek, M. N. (2019). On again, off again: Acute stress response and negative feedback together predict resilience to experimental challenges. Functional Ecology, 33 (4), 619–628.

Zwaan, D. R. de, & Martin, K. (2020). Hierarchical fear: Parental behaviour and corticos-terone release mediate nestling growth in response to predation risk. bioRxiv.

